# *Toxoplasma* TgATG2 works with TgATG9 and TgProp1 to drive autophagic flux for parasite extracellular survival and persistence

**DOI:** 10.64898/2026.07.16.739079

**Authors:** Silvia Masci, Pariyamon Thaprawat, Federica Piro, Tracey L. Schultz, Vern B. Carruthers, Manlio Di Cristina

## Abstract

Chronic infection by *Toxoplasma gondii* depends on long-term survival of bradyzoites within tissue cysts, a parasite stage highly resistant to current therapies and a major barrier to eradication. Autophagy has emerged as critical pathway for bradyzoite persistence, yet the core machinery driving autophagosome formation in *T. gondii* remains poorly defined. Here, we identify TGME49_304630 as TgATG2, a previously uncharacterized, unusually large ATG2-like protein with conserved structural features of lipid-transfer factors. TgATG2 associates with TgATG9 and TgPROP1, key components of the parasite autophagy pathway, supporting its role in a membrane expansion complex required for autophagosome biogenesis. Using independent genetic knockouts, we show that TgATG2 is dispensable for intracellular tachyzoite replication but required for parasite fitness during extracellular stress and, most critically, for bradyzoite autophagy and viability. TgATG2 ablation disrupts autophagic activity in bradyzoites, causing progressive loss of viability and compromised cyst integrity. To overcome limitations of previous indirect assays, we developed a bradyzoite-specific dual-fluorescence TgATG8 reporter that quantitatively measures autophagic flux in *T. gondii* and confirmed TgATG2 as a major contributor. Importantly, TgATG2-deficient parasites are severely impaired during chronic infection in mice, with reduced brain cyst burdens, abnormal cyst morphology, and markedly diminished *ex vivo* bradyzoite viability. Together, these findings establish TgATG2 as a central component of the *T. gondii* autophagy machinery, demonstrate that autophagosome biogenesis is critical for parasite persistence *in vivo*, and define a molecular vulnerability and quantitative platform for targeting autophagy-dependent parasite survival.

**IMPORTANCE:** Current treatments for toxoplasmosis control acute infection but do not eliminate the long-lived tissue cysts responsible for chronic infection. This persistent stage is clinically important because cysts can reactivate in immunocompromised individuals and may also contribute to long-term disease outcomes. A major barrier to developing cyst-targeting therapies is the limited understanding of how bradyzoites maintain viability for extended periods inside host tissues. This study identifies autophagosome biogenesis as a critical survival process in bradyzoites and defines TgATG2 as a key parasite factor required for this pathway. By linking TgATG2-dependent autophagy to cyst viability and persistence *in vivo*, our work highlights parasite autophagy as a potential target for eliminating chronic infection. The autophagic flux reporter developed here also provides an important tool for future studies and for screening approaches aimed at discovering inhibitors of bradyzoite survival.

## INTRODUCTION

Autophagy is a pathway conserved among eukaryotes for delivery of cytoplasmic materials to a cell’s digestive organelle such as the mammalian lysosome or yeast vacuole (1, 2). To maintain homeostasis, cells rely on catabolic pathways that remove old or damaged components to be replaced with new ones through biosynthetic pathways (3). This interplay of efficient turnover and biogenesis of materials is essential to support key development and functions of cells. The core autophagy machinery comprised of autophagy-related proteins (Atg/ATGs) have been extensively studied in mammalian and yeast model systems (4–6).

Macroautophagy delivers cargo to lysosomes via autophagosomes, transient double-membrane organelles generated *de novo* during each degradative cycle. Autophagosome formation depends on autophagy-related proteins, including the Atg2–Atg18–Atg9 complex, which drives phagophore initiation and expansion at the phagophore assembly site (PAS) (7–9). Although multiple organelles may contribute to membrane, lipid delivery from the ER appears central.

Atg2 is a large, rod-shaped lipid transferase that functions at ER–phagophore contact sites. Despite sequence divergence, Atg2 proteins contain conserved N-and C-terminal chorein domains, also found in VPS13 proteins (10–12). These regions mediate membrane binding and tethering: the N-terminal chorein_N segment caps a tubular hydrophobic cavity that transports lipids between membranes, while the C-terminal amphipathic helix promotes membrane association and helps recruit the Atg2–Atg18 complex to the PAS through Atg18 binding to phosphatidylinositol 3-phosphate (PI3P) (13, 14). Together, Atg2 and Atg18 tether the growing phagophore to the ER and support lipid transfer for membrane expansion. Atg9, the only transmembrane core autophagy protein, localizes to ER-and Golgi-derived compartments and interacts with Atg2-containing vesicles to restrict Atg2 to phagophore edges and promote Atg18 association (15). Proper assembly of the Atg9–Atg2–Atg18 complex is therefore essential for establishing ER–phagophore contact sites; disrupting these interactions mislocalizes Atg2 and impairs autophagy (13). Atg2 also functionally cooperates with ER-resident lipid scramblases VMP1 and TMEM41B. While Atg2 transfers lipids from the ER to the phagophore, VMP1 and TMEM41B equilibrate phospholipids across the ER bilayer, maintaining a lipid pool available for Atg2-dependent transport (16). Loss of VMP1 or TMEM41B produces autophagy defects like Atg2 depletion, including impaired phagophore expansion and reduced autophagic flux. Thus, Atg2, VMP1 and TMEM41B likely form a conserved lipid mobilization pathway that supplies membrane for autophagosome biogenesis (17). Despite advances in model eukaryotes, it remains unclear whether Atg2-dependent autophagosome biogenesis is conserved in early-branching eukaryotes and intracellular parasites. This question is especially relevant in Apicomplexa, obligate intracellular parasites with reduced and divergent autophagy-related machinery adapted to parasitism. *Toxoplasma gondii* is a useful model for examining how conserved autophagy pathways have been modified to support parasite survival, differentiation, and persistence. In *T. gondii*, autophagy-related processes contribute to stress adaptation, organelle homeostasis, apicoplast maintenance, and differentiation from tachyzoites to latent bradyzoites, yet the molecular machinery underlying autophagosome biogenesis remains incompletely defined (18–30).

Defining canonical autophagy in Apicomplexa is challenging because many ATG components are absent or highly divergent. Although ubiquitin-like conjugation systems required for ATG8 membrane association are present, several initiation and membrane-expansion factors lack clear homologs, raising debate over whether early-branching eukaryotes possess a canonical autophagy pathway (31). Moreover, some ATG proteins in apicomplexans function in processes unrelated to autophagy, such as apicoplast maintenance (18, 31–33). Nevertheless, autophagic vesicles are induced by nutrient deprivation, cellular stress, or drug treatment in pathogenic parasites, including *T. gondii* and *Plasmodium* spp. (30, 34, 35).

Autophagy regulation also appears divergent in apicomplexans. Homologs of mTOR complexes have not been identified in *Plasmodium* spp., while *T. gondii* encodes a putative TgTOR, though evidence that it inhibits autophagy under nutrient-rich conditions is limited (30, 36). AMPK homologs are present in both *Plasmodium spp.* and *T. gondii*; in *T. gondii*, AMPKα regulates metabolism and parasite growth, but its role in autophagy remains unknown (37, 38). Atg1/ULK1-like kinases have been identified, although they vary substantially outside the conserved catalytic domain (39, 40). By contrast, several initiation-complex components, including ATG13, ATG17, ATG29, and ATG31, lack clear homologs in *Plasmodium* spp. and *T. gondii*. Putative ATG11, FIP200, and ATG101 homologs have been reported in *Plasmodium* spp., but their roles in autophagy remain unresolved (40). These absences suggest that apicomplexans may use alternative mechanisms to initiate autophagosome formation, though whether this reflects ancestral absence or secondary loss remains unclear.

Despite this reduced repertoire, *T. gondii* has a functional autophagy pathway. Tachyzoites and bradyzoites rely on autophagy for survival under extracellular stress and for chronic infection, respectively (24, 26, 41, 42). The putative phospholipid scramblase TgATG9 and its interactor TgPROPIN1, the yeast Atg18 homolog, are required for autophagosome biogenesis and bradyzoite survival (41–43). Autophagic cargo is delivered to the plant-like vacuolar compartment, or PLVAC, a lysosome-like organelle containing hydrolytic enzymes, including five papain-like cathepsins (44, 45). TgCPL, the dominant PLVAC cathepsin L protease, digests host-and autophagy-derived cargo (24, 45–47), and TgCPL-deficient bradyzoites accumulate undigested material in the PLVAC (24). TgATG8 and TgATG9 colocalize with the PLVAC during stress in tachyzoites (30, 48) and in bradyzoites, which exhibit basal autophagy required for cyst persistence. Because TgCPL mediates terminal autophagic degradation, chemical inhibition of TgCPL is commonly used to disrupt and assess autophagic flux in *T. gondii* (49).

Thus, although autophagy is essential for *T. gondii* survival, key steps in autophagosome biogenesis, particularly membrane expansion, remain poorly characterized. We previously showed that TgATG9 is required for autophagosome formation, likely through phospholipid scramblase activity, as it rescues autophagy in yeast Atg9-deficient cells (42). However, despite the conserved interaction between ATG9 and ATG2 in other systems, a putative TgATG2 has not yet been identified.

Herein, we sought to identify and characterize the role of a putative TgATG2 in parasite autophagy using both *in vitro* and *in vivo* experiments. We developed a new autophagic flux assay based on dual-fluorescence tagging of TgATG8 that releases an internal control validated with chemical inhibitors and an autophagy-defective mutant of *T. gondii*. Through colocalization and interaction studies, we provide insights into the putative membrane expansion complex required for *T. gondii* autophagy.

## RESULTS

### Identification of the putative homologue of Atg2 in *T. gondii*

*T. gondii* relies on autophagy for survival under stress in extracellular acute-stage tachyzoites or for persistence of chronic-stage bradyzoites (24, 26, 41). Despite evidence that autophagy is important for parasite survival, the molecular details of this pathway are not well characterized in *T. gondii* (18, 21, 32). Specifically, the early steps of autophagosome biogenesis, including the membrane expansion complex, have yet to be discovered. We previously reported that TgATG9 is required for autophagosome biogenesis, likely through phospholipid scramblase activity due to its ability to partially rescue autophagy in a yeast Atg9 knockout (42). In another study, we showed that the protein TgPROP1, an orthologue of the yeast Atg18, is critical for autophagy and parasite viability during chronic infection (43). Although it is well known that both Atg9 and Atg18/PROP1 interact with Atg2, a putative TgATG2 has yet to be identified. To fill this gap, we interrogated ToxoDB (50) to identify protein candidates displaying similarities to the conserved domains of ATG2 proteins from other organisms. Through this search, we identified a putative TgATG2 candidate (TGME49_304630, annotated as THO complex subunit 2). While TGME49_304630 shared <24% sequence identity with *S. cerevisiae* ScAtg2 or *H. sapiens* HsATG2 (Fig. 1A), it possesses a lipid-transfer-like domain on the N-terminus that has been shown to be important for binding phospholipids near a hydrophobic cavity in model systems (Fig. 1B). TGME49_304630 also contains an ATG2 C-terminal domain (PF09333) that is conserved among ATG2 proteins (Fig. 1B). Based on these features we hereafter refer to this protein as TgATG2. Interestingly, TgATG2 is approximately five times larger than its counterpart in yeast. We used AlphaFold to predict the structure of the N-terminal lipid-transfer-like domain (amino acids 1–1906), which showed a rod-like structure composed of repeated β-sheets, like other ATG2 proteins (Fig. 1C).

**Figure 1.**
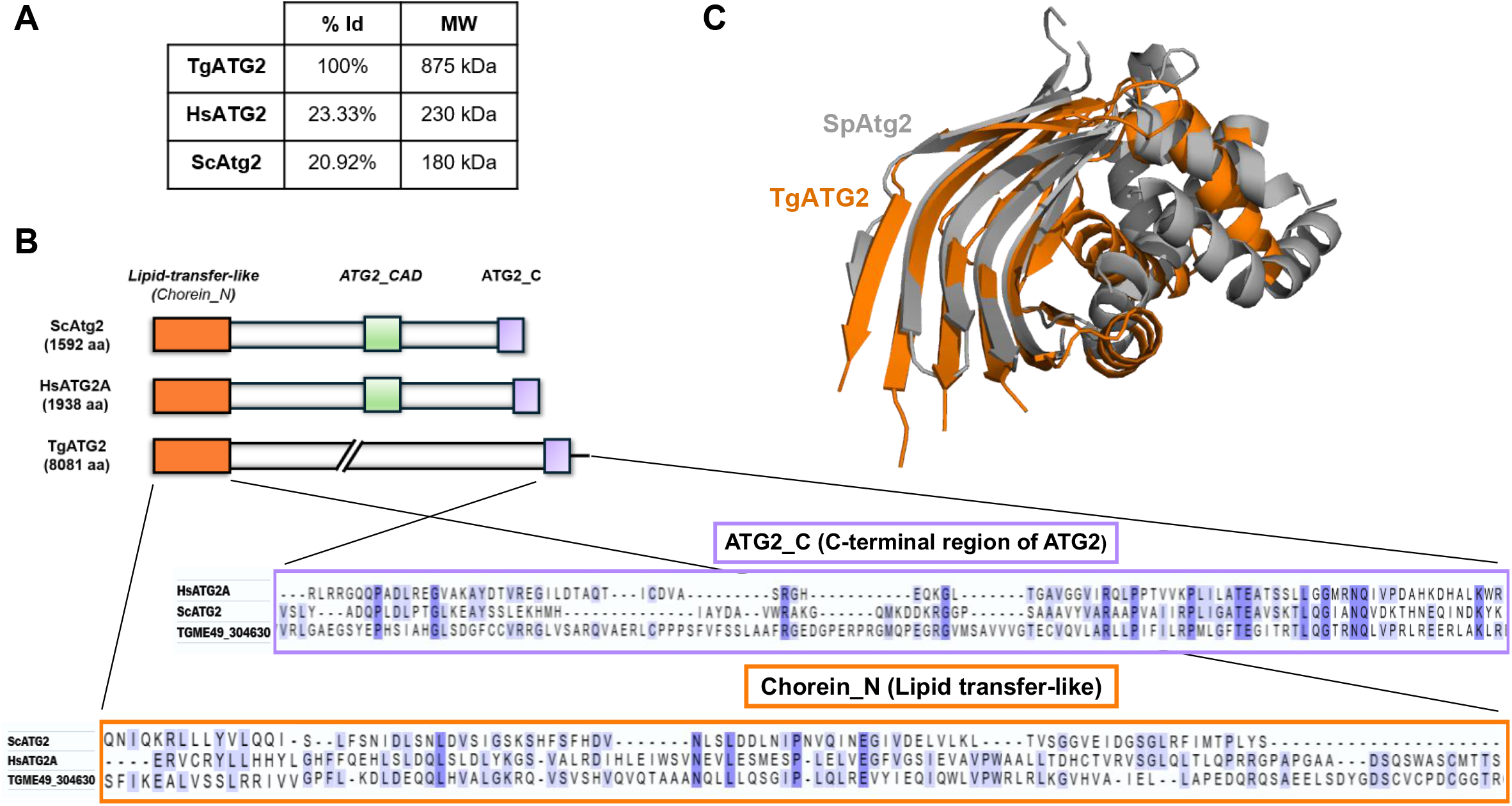
Identification of a putative TgATG2. (**A**) Comparison of amino acid sequence percent identities between TgATG2, human HsATG2, and yeast ScAtg2. Molecular weights for each protein are shown. (**B**) Schematic of ScAtg2, HsATG2A and TgATG2 protein domains with similarities in the lipid-transfer-like and ATG2 C-terminal domains. (**C**) Predicted structure of TgATG2 (amino acids 183-664, sans unstructured regions) with a fold aligned to the corresponding region of *Schizosaccharomyces pombe* Atg2 (6A9J), forming a hydrophobic groove for lipid transfer.

### TgATG2 interacts with TgATG9

To determine whether TgATG2 might function together with TgATG9 in bradyzoite autophagy, we performed colocalization and protein–protein interaction analyses of the two proteins. Since autophagy proteins such as TgATG9 are expressed at low abundance in *T. gondii*, we generated a double-tagged transgenic parasite line (ME49ku80hxg/DD-BFD1/TgATG2-smHA/TgATG9-smMyc) in a background that allows for a greater density of tissue culture infection with conditional stabilization of a master regulator of bradyzoite differentiation (DD-BFD1 stabilized with addition of Shield-1) (51). The spaghetti monster (“sm”) tags appended to the C-terminus of TgATG2 or TgATG9 contain 10 copies of their respective epitopes to allow for higher sensitivity in detection of proteins (52, 53) (Fig. 2A). Immunofluorescence staining of *in vitro* cysts containing bradyzoites showed significant colocalization between TgATG9 and TgATG2 relative to the TgATRX control (an apicoplast marker) (Fig. 2B and 2C). Given that TgATG9 and TgATG2 appeared to colocalize within bradyzoites, we wanted to further investigate whether the two proteins interact. We performed co-immunoprecipitation followed by western blot analysis using TgATG2 as the bait protein from total bradyzoite lysates (Fig. 2D). TgATG9 co-immunoprecipitated with TgATG2, and this interaction was further validated by performing co-immunoprecipitation followed by dot blot to allow the detection of the high-molecular-weight TgATG2 bait protein (Fig. 2E). Together, these findings indicate that TgATG2 colocalizes and interacts with TgATG9, warranting continued investigation into the role of TgATG2 in *T. gondii* autophagy.

**Figure 2.**
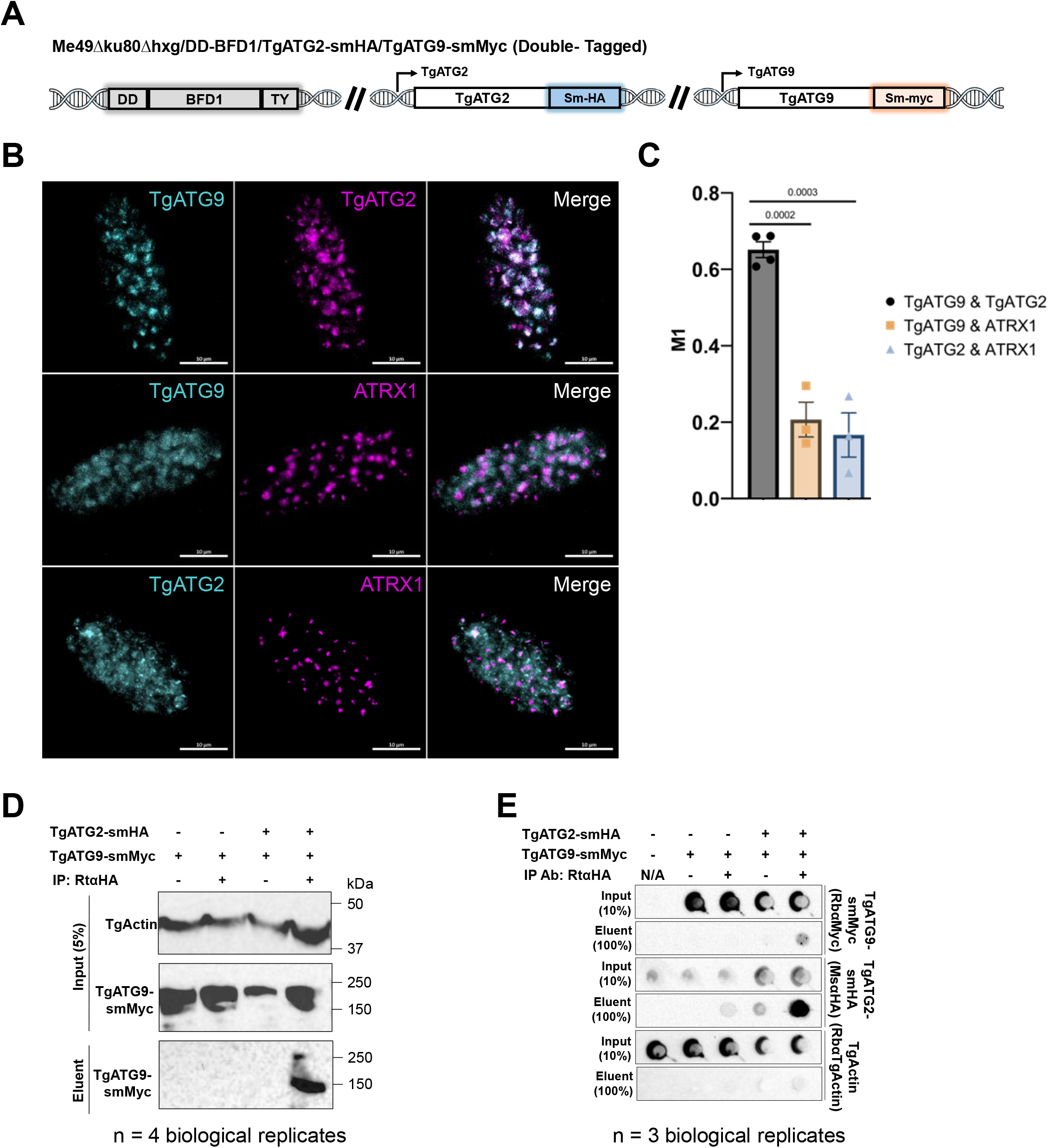
TgATG2 interacts with TgATG9. **A)** Schematic showing endogenous double-tagging of TgATG2 and TgATG9 with spaghetti-monster-HA (smHA) and spaghetti-monster-Myc (smMyc), respectively. **B)** Immunofluorescence staining of in vitro differentiated cysts for colocalization of TgATG9-smMyc and TgATG2-smHA along with the apicoplast marker ATRX1 used as a control. Scale bars, 10μm. **C)** Quantification of M1 colocalization coefficients using CellProfiler software. Bars represent mean ± SD. Unpaired t-test (n=3-4 biological replicates). **D)** Western blot of TgATG2 and TgATG9 co-immunoprecipitation (coIP) using anti-HA IP antibody to capture TgATG2 as the bait protein. TgATG9 is only enriched in the coIP eluent from TgATG2-smHA/TgATG9-smMyc double-tagged parasites compared to non-IP antibody or TgATG9-smMyc single-tagged controls. TgATG2 is not visualized on blot due its high molecular weight. TgActin was used as a loading control. Representative blot shown (n = 4 biological replicates). **E)** Dot blot of TgATG2-smHA and TgATG9-smMyc coIP using anti-HA IP antibody to capture TgATG2 as the bait protein. TgATG9 is only enriched in the coIP eluent from TgATG2-smHA/TgATG9-smMyc double-tagged parasites compared to untagged, non-IP antibody or TgATG9-smMyc single-tagged controls. Order of blot (rows) shows the order the dot blot was probed and stripped for re-probing. TgActin was used as a loading control. Representative blot shown (n = 3 biological replicates).

### TgATG2 also interacts with TgPROP1

In other organisms, ATG2 interacts not only with ATG9 but also with ATG18. Two PROPPINs (TgPROP1 and TgPROP2) have been identified in *T. gondii* based on homology-directed searches for WD40 domain-containing proteins with similarities to *S. cerevisiae* Atg18. In a recent study, we showed that TgPROP1 is critical for bradyzoite autophagy, likely serving as the homologue of the yeast ATG18 (43). Since our TgATG2 candidate interacts with TgATG9, we would expect an interaction of this protein with TgPROP1 as well. We first assessed whether TgATG2 and TgPROP1 colocalization by immunofluorescence analysis. To perform colocalization and interaction analyses of TgATG2 and TgPROP1, we generated a double-tagged transgenic parasite line (ME49ku80hxg/DD-BFD1/TgATG2-smMyc/TgPROP1-smHA) (Fig.3A).

**Figure 3.**
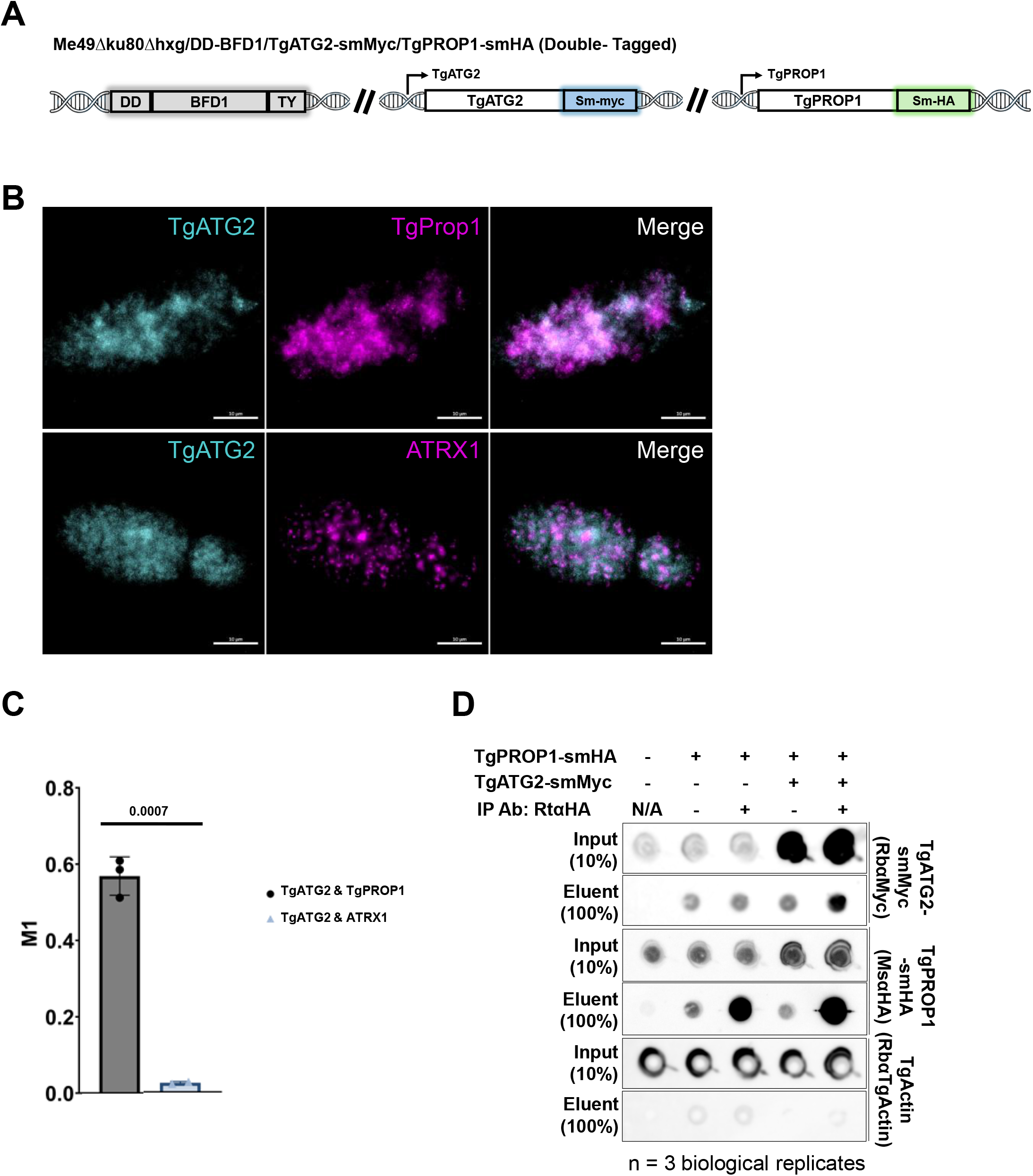
TgProp1 interacts with TgATG2. **A)** Schematic showing endogenous double-tagging of TgATG2 and TgProp1 with spaghetti-monster-Myc (smMyc) and spaghetti-monster-HA (smHA), respectively. **B)** Immunofluorescence staining of in vitro differentiated cysts for colocalization of TgProp1-smHA and TgATG2-smMyc along with the apicoplast markerATRX1 used as control. Scale bars, 10μm. **C)** Quantification of M1 colocalization coefficients using CellProfiler software. Bars represent mean ± SD. Unpaired t-test with p-values (n=2-3 biological replicates). **D)** Dot blot of TgATG2 and TgProp1 co-immunoprecipitation (coIP) using anti-HA IP antibody to capture TgProp1 as the bait protein. TgATG2 is only enriched in the coIP eluent from TgProp1-smHA/TgATG2-smMyc double-tagged parasites compared to non-IP antibody or TgProp1-smHA single-tagged controls. Order of blot (rows) shows the order the dot blot was probed and stripped for re-probing. TgActin was used as a loading control. Representative blot shown (n = 3 biological replicates).

Immunofluorescence staining of *in vitro* cysts containing bradyzoites showed significant colocalization between TgATG2 and TgPROP1 relative to the TgATRX control (Fig. 3B and 3C). Next, we performed co-immunoprecipitation followed by dot blot analysis using TgPROP1 as the bait protein from total bradyzoite lysates and found that TgATG2 co-immunoprecipitated with TgPROP1 (Fig. 3D). Collectively, our findings suggest that TgATG2 likely works in conjunction with TgATG9 and TgPROP1 in a putative membrane expansion complex.

### TgATG2 is important for parasite autophagy

Since TgATG2 was found to directly interact with TgATG9 and TgPROP1, we aimed to determine its importance in bradyzoite autophagy. Because the ME49ku80hxg/DD-BFD1 used above doesn’t readily form brain cysts, we instead used ME49Δku80Δhxg, which is highly cystogenic in mice. We first appended an sm-HA tag to the C-terminus of TgATG2 in ME49Δku80Δhxg to confirm the expression of this protein in this strain (Fig. 4A). Immunofluorescence analysis of tagged tachyzoites showed low expression of TgATG2 that colocalized with the ER marker SERCA (Fig. 4B). To determine its importance in autophagy, we generated a stable genetic knockout of TgATG2 using CRISPR-Cas9 by replacing the entire genomic sequence with the chloramphenicol acetyltransferase (CAT) selectable marker (54, 55) (Fig. S1). Due to the size of TgATG2, we were unable to generate a genetic complement of the TgATG2 knockout (KO). Therefore, we generated an independent stable knockout of TgATG2 using different CRISPR-Cas9 guide RNAs by replacing the N-terminal half of the genomic sequence with the hypoxanthine-xanthine-guanine phosphoribosyltransferase (HXGPRT) selectable marker (MΔTgATG2′) (Fig. S1). Previous work showed that starved extracellular tachyzoites activate autophagy to counteract this stress, and inhibition of this process through ablation of the TgATG9 protein reduced the extracellular survival of tachyzoites (26). To assess whether TgATG2 ablation had the same effect as TgATG9 on tachyzoite survival during extracellular exposure, we mechanically extruded tachyzoites of wild type and both TgATG2 KOs from host cells and incubated parasites in Hanks’ Balanced Salt Solution (HBSS) for different times of exposure to extracellular stress. The lack of TgATG2 resulted in a reduction of extracellular tachyzoite survival after 30 and 60 minutes of incubation in HBSS (Fig. 5A). The invasion defect (Fig. 5B) and the reduced number of plaques formed in plaque assays (Fig. 5C and 5D) observed for both TgATG2 KO strains were consistent with a decrease in extracellular tachyzoite viability.

**Figure 4.**
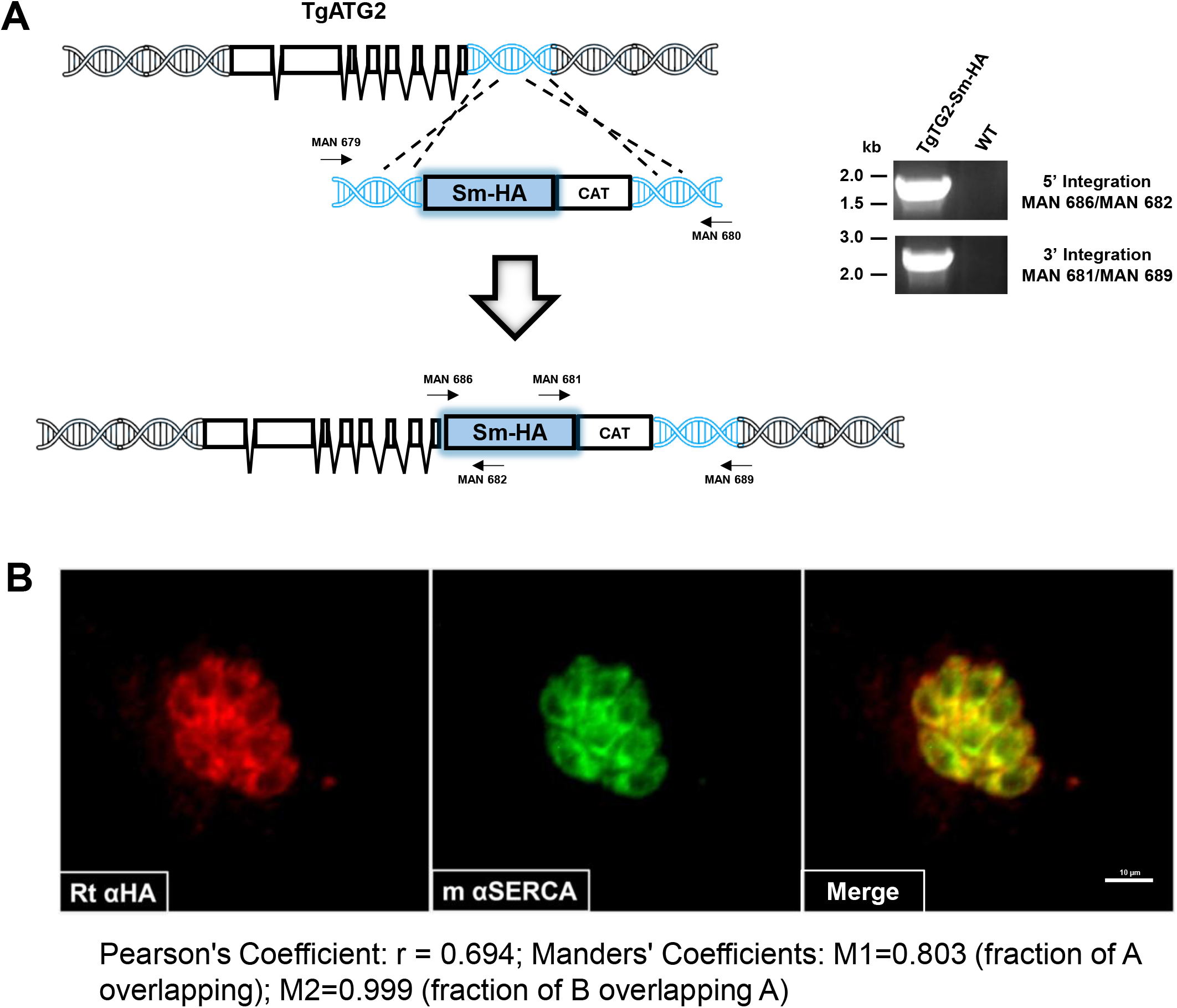
TgATG2 is an ER protein. **A)** Schematic representation of the strategy to tag TgATG2 at its C-terminus with the Spaghetti monster HA tag (Sm-HA). Right panel: PCR validation to confirm the successful insertion of the Sm-HA coding sequence at the 3’ of the TgATG2 gene in the Me49Δku80 strain (hereafter Me49). All primer sequences and gRNAs are reported in Table S1. **B)** Representative images of parasites processed for IFA using rat anti-HA (red) and mouse anti-TgSERCA (green) antibodies. TgATG2 and TgSERCA co-localization indicated that TgATG2 is resident on the endoplasmic reticulum (Scale bar, 10μm).

**Figure 5.**
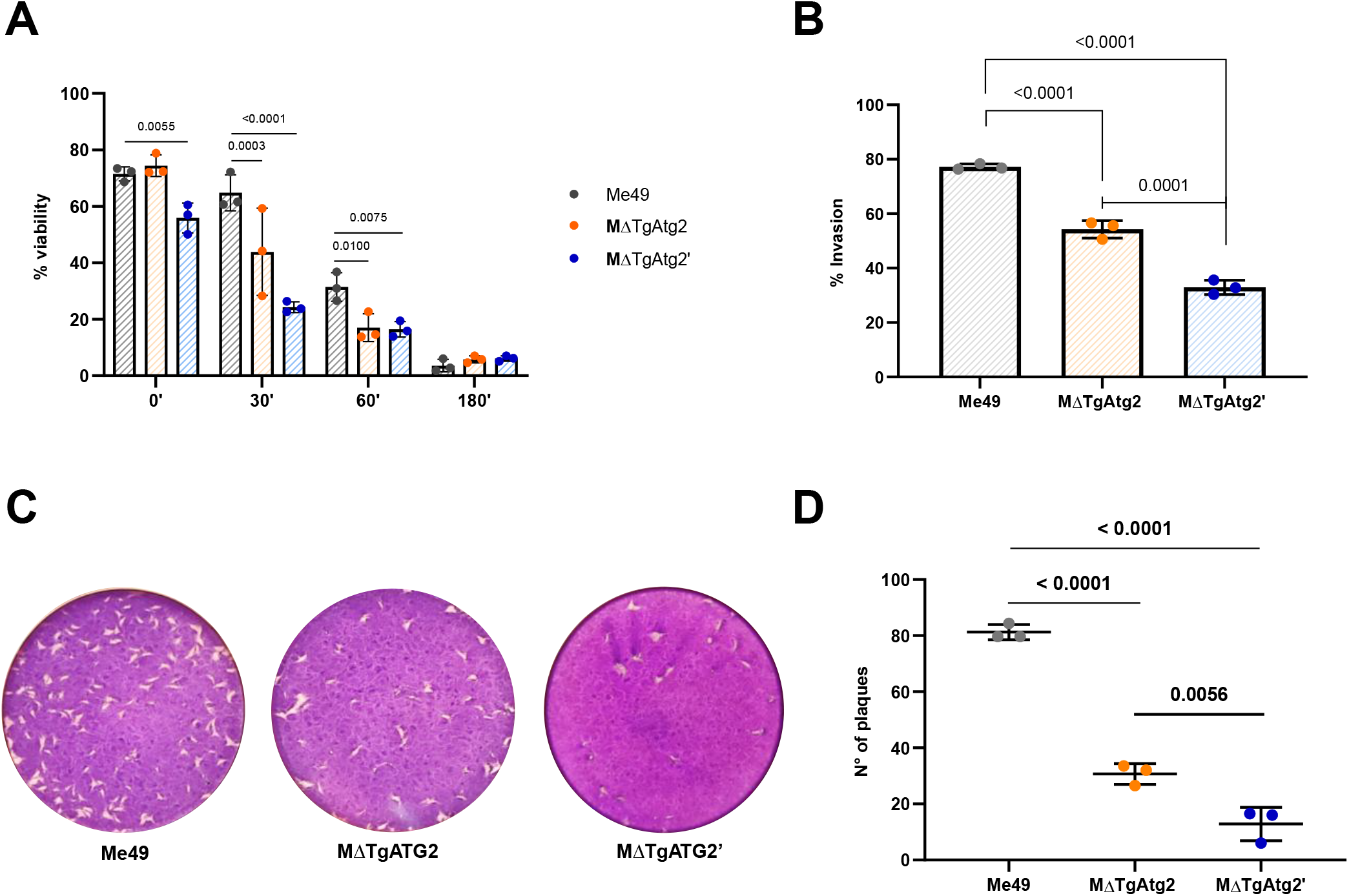
TgATG2 is important for extracellular survival and invasion. **A)** PMA viability assay of WT and ΔTgATG2 extracellular tachyzoites mechanically extruded from HFF and incubated in HBSS for the indicated time points. Two-way ANOVA with Dunnett’s multiple comparisons was performed. **B)** TgATG2 ablation slightly impacts host cell invasion. Shown are the results of a red-green invasion assay of tachyzoites after 20 min of incubation with HFF cells. Parasites were stained as described in Materials and Methods. The graph represents means ± SD from three independent experiments. **C)** and **D)** Deletion of TgATG2 resulted in a significant reduction of the number of plaques formed compared to WT, as assessed by plaque formation after 12 days. One-way ANOVA with Tukey’s multiple comparisons was performed in **(B)** and **(D)** analyses.

While autophagy is not required during intracellular tachyzoite growth, it becomes essential for bradyzoite maintenance within cysts. Therefore, we assessed autophagic function of the TgATG2 KOs by differentiating *in vitro* bradyzoites for 7 days, followed by treatment with LHVS or DMSO for 3 days prior to CytoID staining, which selectively stains autolysosomes (24, 41–43, 56, 57). LHVS inhibits the major cathepsin protease (TgCPL) within the parasite digestive organelle, the plant-like vacuolar compartment (PLVAC) (49). TgATG2 KO parasites were unable to accumulate Cyto-ID positive structures after LHVS treatment compared to WT (Fig. 6). If TgATG2 contributes to autophagy like TgATG9 and TgPROP1 then we reasoned that TgATG2 deficient bradyzoites should show reduced viability. To test this, we differentiated *in vitro* bradyzoites of WT and TgATG2 KOs, then assessed the viability of bradyzoites after 7 and 14 days post-differentiation using the PMA bradyzoite viability assay (58). We observed a drop in bradyzoite viability of about 40% and 95% in ΔTgATG2 parasites after 7 and 14 days post-conversion, respectively (Fig. 7). These results provide evidence for the critical role of TgATG2 in bradyzoite autophagy and further confirm the crucial role played by autophagy in the bradyzoite stage.

**Figure 6.**
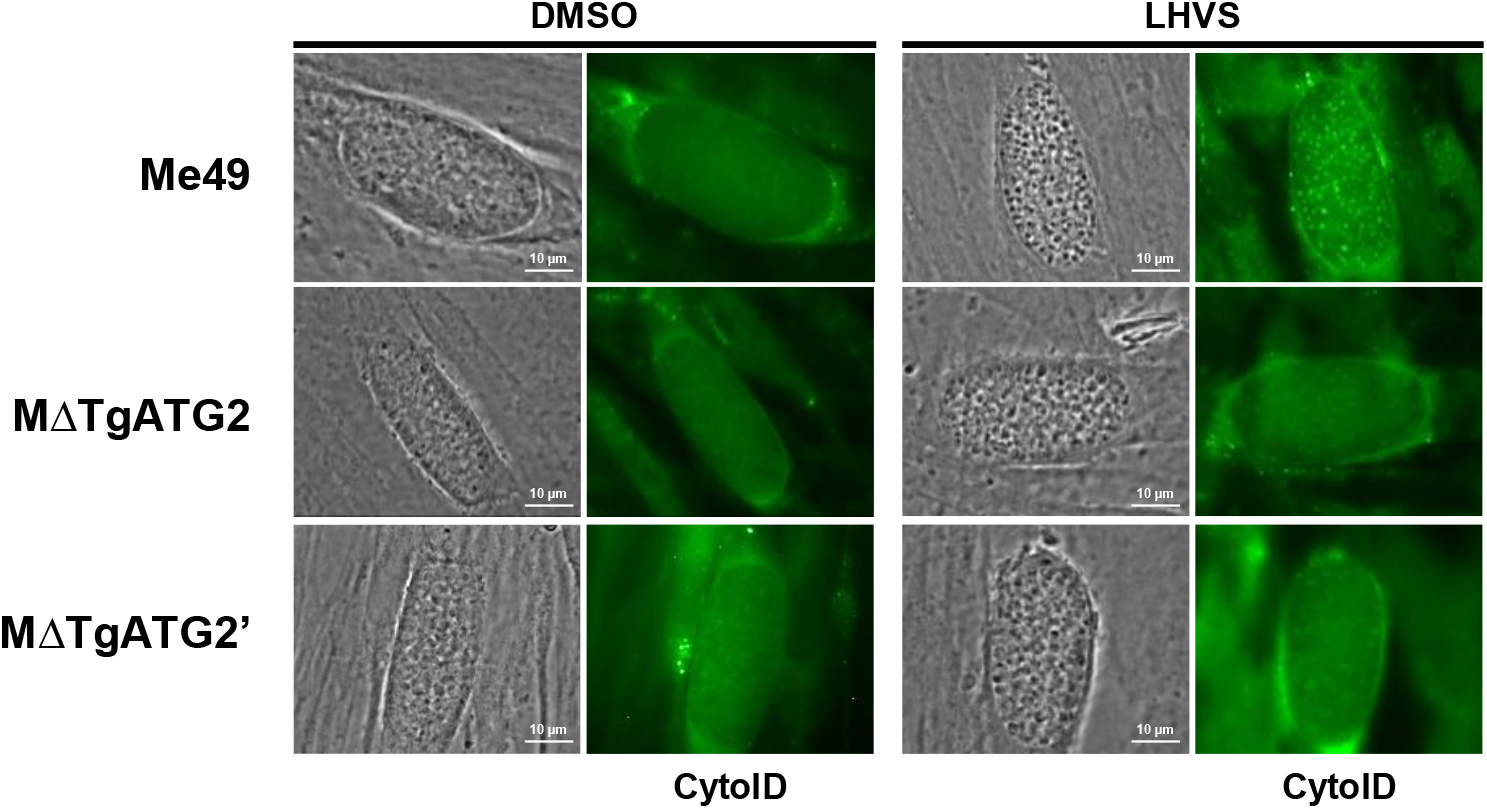
Bradyzoites lacking TgATG2 show diminished staining of autolysosomes. Autolysosome staining (CytoID) of in vitro Me49, MΔTgATG2 or MΔTgATG2’ differentiated cysts treated with vehicle control (DMSO) or LHVS for 72 h (n=3 biological replicates, representative images shown). Scale bars, 10 μm.

**Figure 7.**
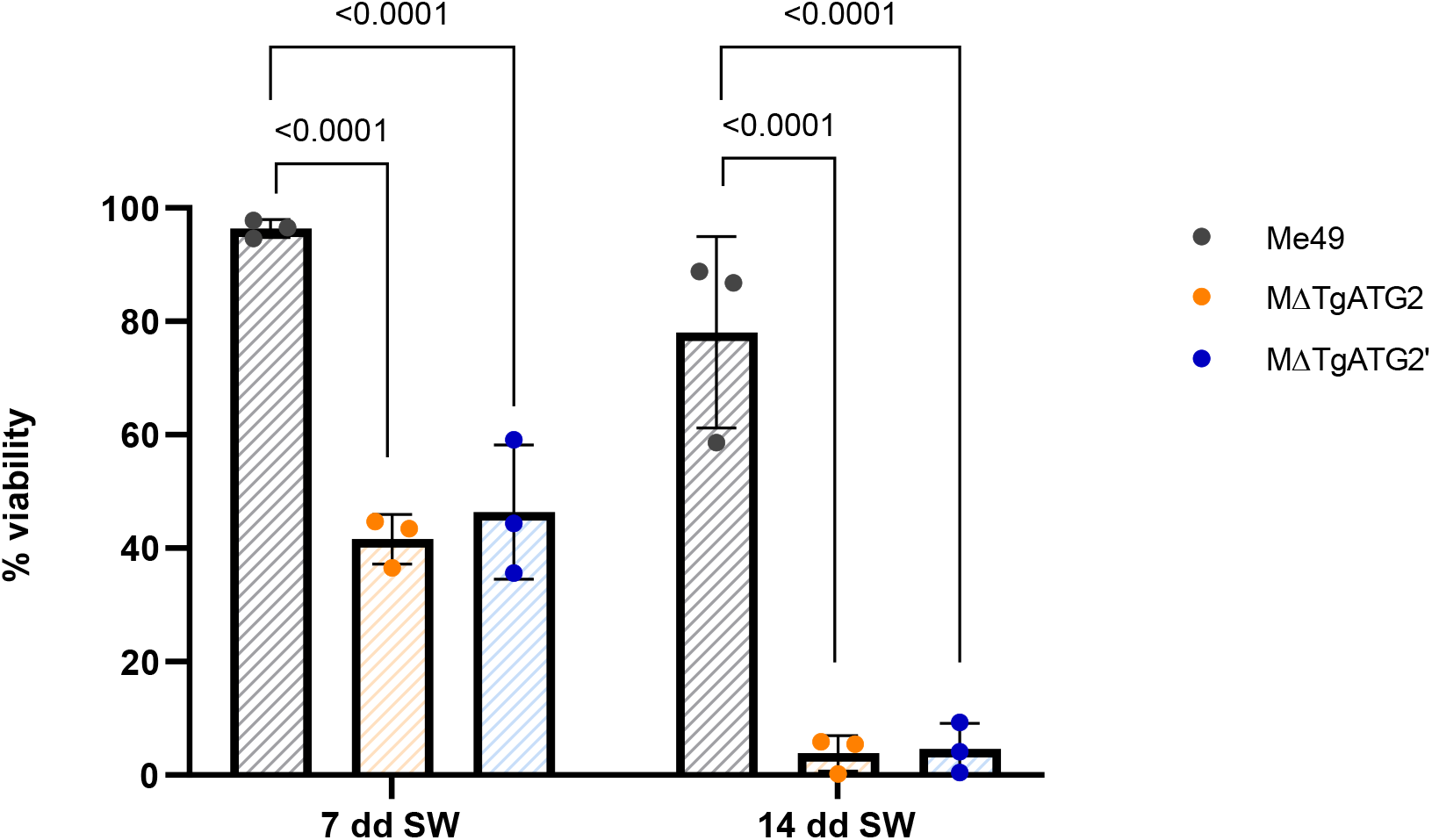
Bradyzoites lacking TgATG2 show a time dependent loss of viability. WT, MΔATG2 and MΔATG2’ tachyzoites were converted into bradyzoites by alkaline induction (sw) for either 7 (left panel) or 14 days (right panel) and analyzed for viability using the PMA/qPCR assay. Error bars indicate the SD from three independent experiments. One-way ANOVA with Sidak’s multiple comparisons was performed.

### Development of a stage-specific reporter to measure autophagic flux in *T. gondii*

Although we have extensively used CytoID staining in our studies of bradyzoite autophagy, this assay is not a direct measure of autophagic flux and relies on LHVS treatment for autophagic material to accumulate within autolysosomes visualized by the CytoID dye. Therefore, most reports of bradyzoite autophagy have been limited to observations of accumulated autolysosomes as readouts for autophagic function in *T. gondii* (24, 41–43). To address this limitation, we aimed to develop a bradyzoite-specific autophagic flux reporter that would allow for more robust quantification of parasite autophagic flux. In yeast and mammalian cells, many autophagy reporters rely on the lipidation of Atg8/LC3 onto autophagic membranes and its delivery to the cell’s digestive organelle to indicate flux (59–65). Once delivered to the digestive organelle, Atg8/LC3, along with any fluorescent markers appended to its N-terminus, are degraded within the lumen of the proteolytic compartment (66, 67). Assessments of Atg8/LC3 processing can be determined by western blotting for protein levels or by pH-specific quenching of pH-sensitive fluorescent proteins (such as GFP) (68–70). We modeled our autophagic flux reporter based on the GFP-LC3-RFP-LC3ΔG fluorescent probe, which is cleaved by ATG4 to release RFP-LC3ΔG, as an internal control, and GFP-LC3 (66, 70). We generated a plasmid containing tdTomato-2A-GFP-TgATG8 under the bradyzoite-specific promoter BAG1 (pBAG1) and introduced this construct into ME49Δku80Δhxg parasites at the BAG1 locus for exogenous expression of the autophagic flux reporter (ME49Δku80/tdTomato-2A-GFP-TgATG8, hereafter the strain carrying the autophagic sensor is referred as M/Sn, Fig. S2). The presence of the 2A skip peptide (specifically, T2A from *Thosea asigna* virus 2A) induces ribosome skipping during protein translation (71, 72). This results in the release of tdTomato, which remains cytosolic as an internal control, and GFP-TgATG8 (Fig. 8A). We first validated the autophagic flux reporter by deleting the TgATG9 gene in M/Sn. TgATG9 is known to be essential for a functional autophagy pathway and thus can be used to validate the new autophagic flux reporter. Bradyzoites were differentiated *in vitro* for 7 days prior to fixation and imaging. The relative GFP:tdTomato ratio for a given cyst was calculated. Compared to wild-type cysts, M/SnΔTgATG9 resulted in a significant increase in the GFP:tdTomato ratio, suggesting a decrease in autophagic flux due to inhibition of proteolytic turnover within the PLVAC (Fig. 8B and 8C). Next, we utilized our new reporter to quantify the disruption to bradyzoite autophagy in the TgATG2 KOs compared to WT. There was a significant increase in the GFP:tdTomato ratio in TgATG2 KO parasites compared to WT controls (Fig. 8B and 8C).

**Figure 8.**
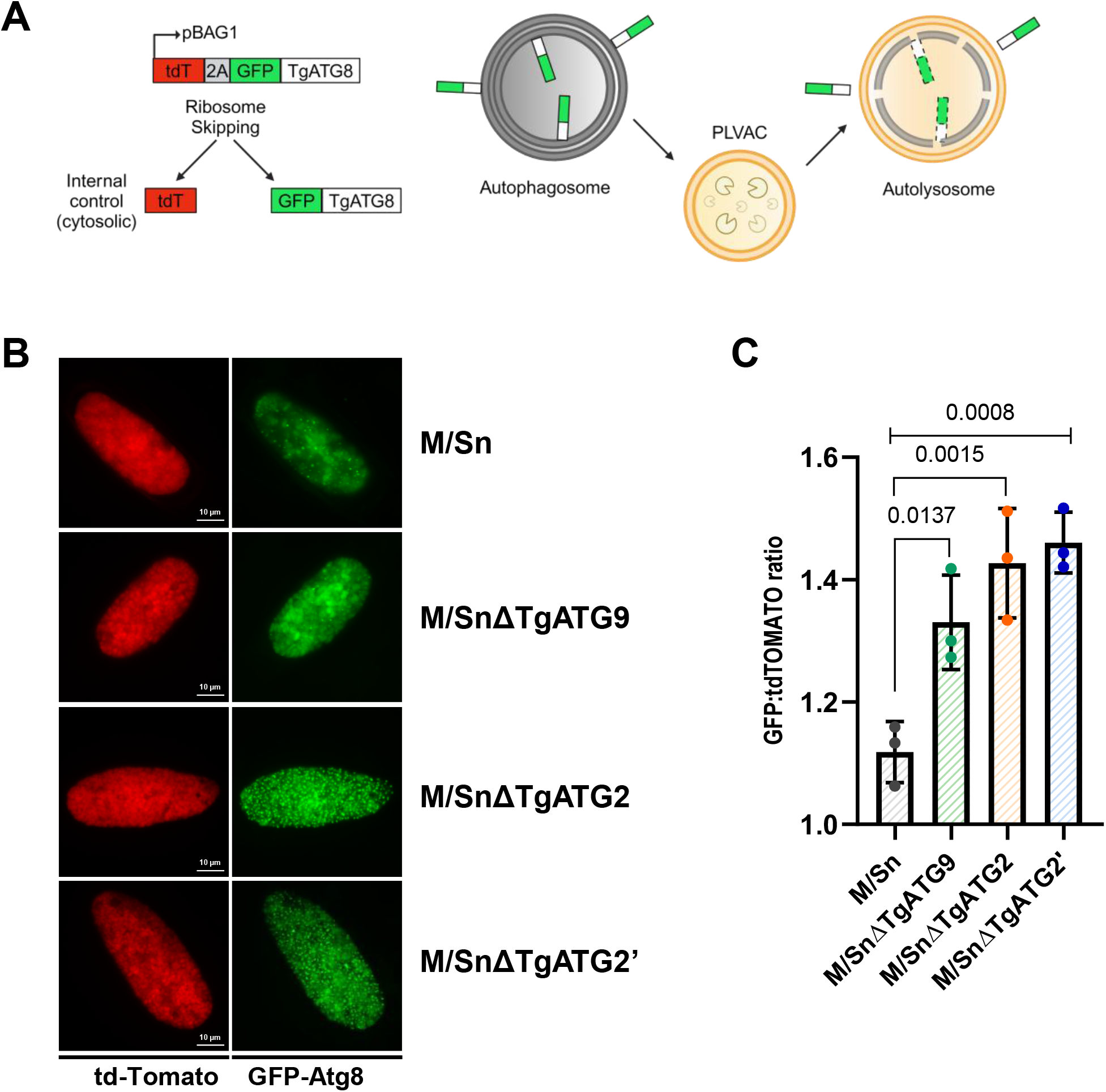
Development of a bradyzoite-specific autophagic flux reporter in *T. gondii*. **(A)** An autophagic flux reporter was generated by exogenous expression of TgATG8 with a tdTomato-2A-GFP tag appended on the N-terminus and driven by the bradyzoite-specific BAG1 promoter (pBAG1). The 2A peptide results in ribosome skipping to generate tdTomato (tdT) released as an internal cytosolic control and GFP-TgATG8 that is subsequently lipidated onto autophagic membranes. Upon delivery of autophagosomes to the PLVAC, a fraction of GFP-TgATG8 is degraded, resulting in decreased GFP fluorescent signals that can be quantified and normalized to tdTomato internal controls (GFP:tdTomato ratio). A lower ratio indicates higher autophagic flux. **B)** Bradyzoites lacking TgATG2 have lower autophagic flux. Widefield fluorescence microscopy images of in vitro differentiated reporter cysts WT (Me49Δku80/tdTomato-2A-GFP-TgATG8, hereafter M/Sn), M/SnΔTgATG2 or M/SnΔTgATG2, n = 3 biological replicates with at least 50 cysts imaged per condition, representative images shown. Scale bars, 10 μm. **C)** Quantification of the GFP:tdTomato ratio from **(B)**. Bars represent mean ± SD. Statistical analysis was done using One-way ANOVA with Dunnett’s multiple comparisons, p-values denoted above bars.

As shown in other organisms, such as *Saccharomyces cerevisiae*, GFP fused to Atg8 is cleaved from Atg8 by hydrolytic enzymes when autophagosomes are delivered to yeast vacuoles. Assessing the ratio of free GFP to total GFP, defined as free GFP plus GFP-Atg8, using anti-GFP antibodies in western blot experiments, is used to quantify autophagic flux (59–65). Therefore, we first evaluated whether free GFP was detectable in M/Sn parasites converted into bradyzoites by growth in alkaline medium for 1 week. Since free GFP was barely visible due to rapid degradation by PLVAC proteases, we treated bradyzoite cultures with LHVS for 1 and 3 days to inhibit the CPL protease and delay free GFP degradation (Fig. 9A). Three days of LHVS treatment resulted in greater accumulation of free GFP in western blot analysis, allowing better determination of its quantity to measure basal autophagic flux in bradyzoites (Fig. 9B). Following this pipeline, we measured the autophagic flux of parasites deficient in both TgATG9 and TgATG2. Loss of TgATG9 appeared to completely abolish autophagic flux, as no free GFP was detectable in any of the three biological replicates, whereas both TgATG2 KO strains showed a reduction in autophagic flux of approximately 65–70% (Fig. 9C and 9D).

**Figure 9.**
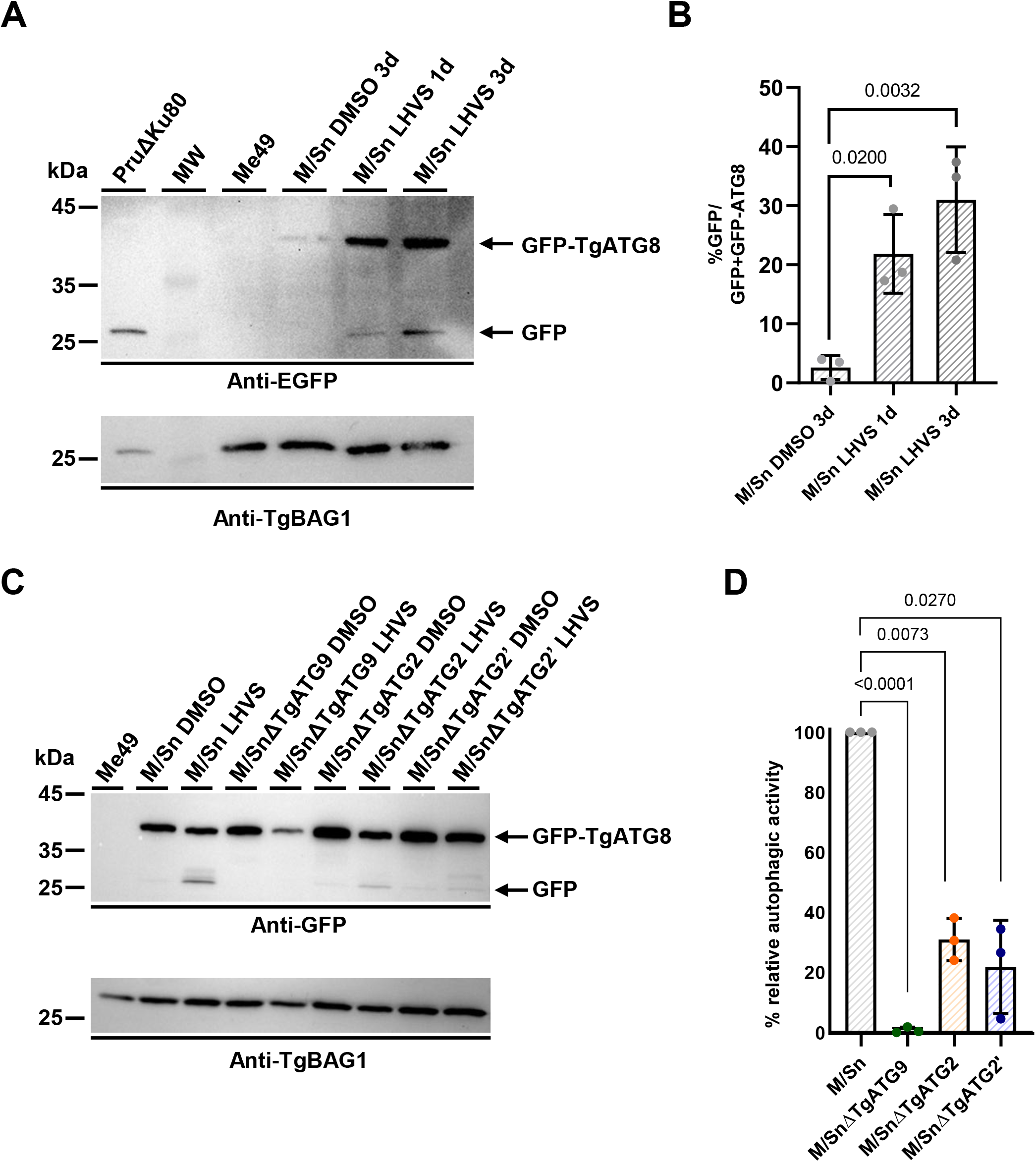
Autophagy activity quantified by Western Blot (WB) assay. **A)** Validation of the WB assay to quantify the autophagy flux of M/Sn bradyzoites. M/Sn tachyzoites were converted into bradyzoites by growth in alkaline media for 7 days and then treated with DMSO (viacle) or LHVS for 1 or 3 days. Bradyzoites were then recovered from cysts by pepsin treatment, lysed in Laemmli Lysis buffer, boiled and ran in 10% SDS-Page gel. After transferring to PVDF membrane, proteins were incubated with mouse anti-GFP antibodies. After secondary HRP antibody incubation, filters were analyzed using a WB detection system. **B)** Quantification of free GFP from 3 biological replicates of WB from **(A)**. One-way ANOVA with Dunnett’s multiple comparisons was performed. Three days of LHVS treatment resulted in higher accumulation of free GFP compared to 1 day treatment. **C)** WB analysis to quantify autophagy flux of ΔTgATG9 and ΔTgATG2 bradyzoites compared to WT bradyzoites (placed as 100% autophagy flux in normal condition). Three biological replicates from **(C)** were used to quantify free GFP and generated a plot to measure the impact of TgATG9 and TgATG2 on the autophagy flux **(D)**. Lack of TgATG9, known to be essential for functional autophagy in *T. gondii* was used as reference of reduced or absent autophagy. Parasites were converted into bradyzoites by growth in alkaline media for 7 days and then treated with DMSO or LHVS for further three days in alkaline media. Samples were processed as in **(A)**. Brown-Forsythe and Welch ANOVA test (One-way ANOVA) with Dunnett’s T3 multiple comparisons was performed.

Thus, we have demonstrated that bradyzoite autophagy can be quantified using our newly developed autophagic flux reporter via genetic ablation of TgATG9 and TgATG2.

### TgATG2 is critical for *T. gondii* persistence in a mouse model of chronic infection

To validate the observed *in vitro* phenotype, we determined the necessity of TgATG2 for persistence of *T. gondii in vivo* during chronic infection. We infected mice with wild type (ME49Δku80Δhxg), TgATG2 KO, or TgATG2′ KO parasites. At 5 weeks post-infection, brains from infected mice were collected and cyst burdens were enumerated. We observed a significant ∼3-fold decreased cyst burden in the brains of mice infected with TgATG2 KO or TgATG2′ KO parasites compared to WT (Fig. 10A). To evaluate the viability of *ex vivo* parasites, we isolated bradyzoites from a subset of the brains harvested (3 per strain) using pepsin treatment followed by plaque assay normalized to the number of parasite genomes by qPCR (73). WT bradyzoites produced quantifiable plaques, while minimal to no plaques were observed for the TgATG2 KO strains (Fig. 10B). We visually examined *ex vivo* cysts from mouse brains via light microscopy and found that while WT cysts appeared healthy with no morphological abnormalities, TgATG2 KO and TgATG2′ KO cysts appeared unhealthy, with less defined cyst walls, bloated bradyzoites, and increased gaps between bradyzoites within cysts (Fig. 10C).

**Figure 10.**
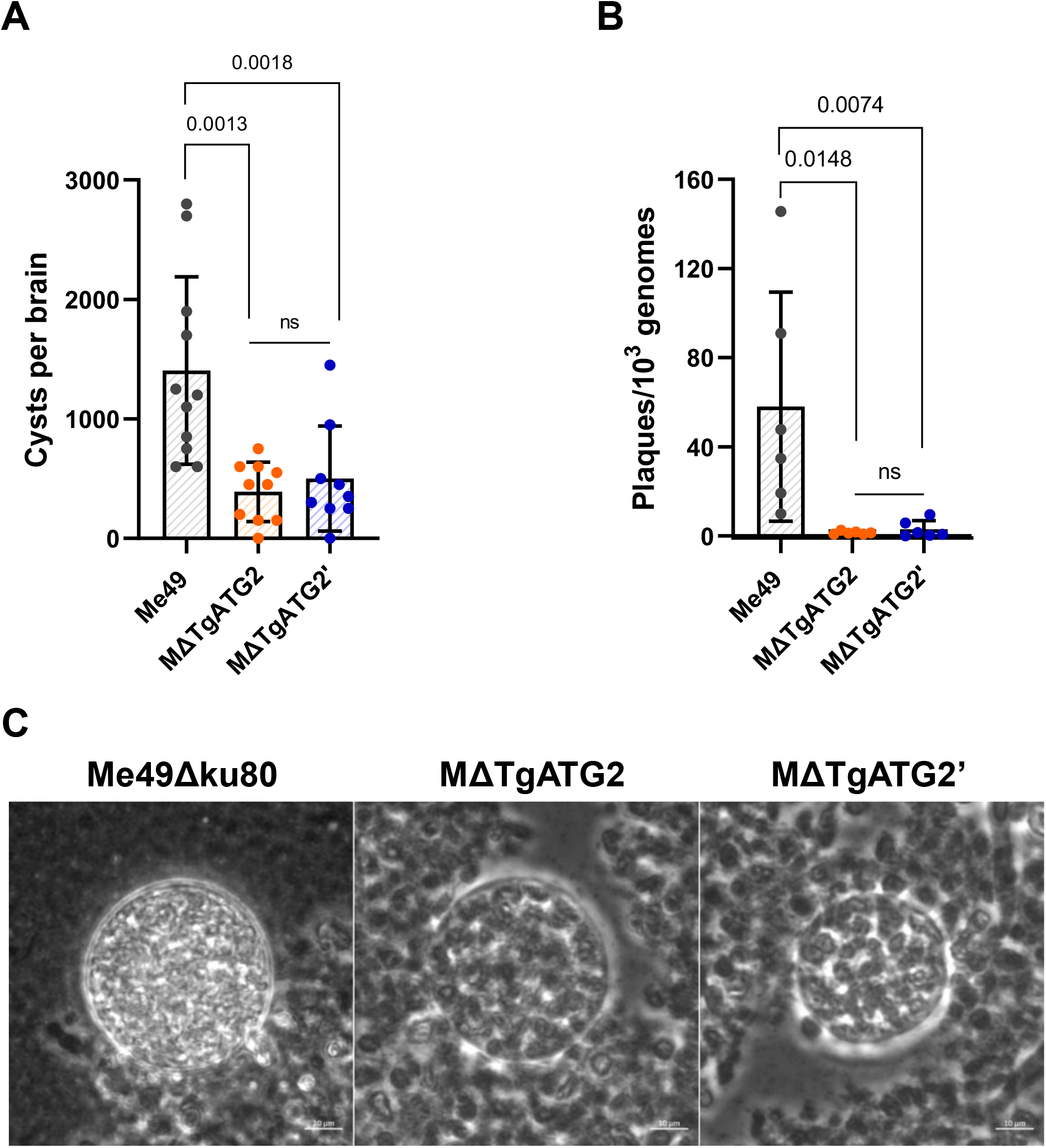
The putative TgATG2 is required for *T. gondii* persistence and viability during chronic infection. **A)** Cyst burdens per brain quantified from chronically infected mice at 5-weeks post-infection with WT, ΔTgATG2 or ΔTgATG2’ parasites. Bars represent mean ± SD. Statistical analysis was done using the Kruskal test Dunn’s multiple comparisons, p-values denoted above bars. **B)** Viability of *ex vivo* bradyzoites harvested from infected mouse brains **(A)** reported as number of plaques normalized to parasite genomes. Bars indicate mean ± SD. Statistical analysis was done using the Kruskal-Wallis test with Dunn’s multiple comparisons, p-values denoted above bars. **C)** Representative images of ex vivo cysts of WT, MΔTgATG2 or MΔTgATG2’ from infected mouse brains under light microscopy.

Thus, our findings confirm the importance of TgATG2 for bradyzoite autophagy and viability of *T. gondii* during chronic infection.

## DISCUSSION

While autophagy is dispensable for *Toxoplasma gondii* tachyzoites during their intracellular life, it becomes necessary when parasites egress from infected host cells and exposed to extracellular stress (26). Autophagy is also an essential process during the chronic phase of infection, characterized by slow-growing bradyzoites contained within tissue cysts. Although the precise reason for autophagy being critical for bradyzoites is still unknown, it could relate to a need for maintaining organellar homeostasis during this latent stage. Whereas with each round of cell division tachyzoites has the potential to divert damaged organelles to the residual body (a membrane bound structure containing organellar remnants formed at the posterior end of the parasite), bradyzoites do not generate a residual body during cell division. Also, damaged organelles that are retained by tachyzoites are diluted out by substantial new biosynthesis during rapid growth, which is likely more limited in bradyzoites and would need to be counterbalanced by the turnover of such organelles by autophagy. Regardless of the underlying basis, autophagy represents a vulnerability of bradyzoites, with its inhibition leading to cyst degeneration and the potential eradication of the chronic phase of *T. gondii*. However, studying autophagy in *T. gondii* has been limited not only by a lack of identified TgATG candidates but also by a lack of robust assays that directly assess autophagic function or flux. Prior studies have relied on indirect measures of autophagic function, such as parasite viability or burden of chronic infection in mouse models (21, 24, 26, 41, 74). The accumulation of undigested autophagic material upon chemical inhibition or genetic ablation of the major cathepsin protease L (TgCPL) in the PLVAC has also served to inform the status of autophagy within bradyzoites (24, 41–43). In this study, we developed a bradyzoite-specific autophagic flux assay based on dual-fluorescence tagging of TgATG8 that releases an internal control, validated with autophagy-defective mutants of *T. gondii*, such as TgATG9 and TgATG2, the latter of which was identified and characterized in this work.

Several ATG proteins have been identified in *T. gondii*, but only TgATG9 and TgPROP1, the Atg18/WIPI orthologue, do not appear to play a dual role in both canonical autophagy and apicoplast maintenance, a plastid organelle housing essential metabolic functions in *T. gondii* (19, 20, 27, 33, 75, 76). Instead, they are key components of the core autophagy machinery and are important for cyst persistence during the chronic phase of infection (26, 41, 43, 74). Given that Atg9 and Atg18 function at the membrane expansion step in cooperation with Atg2 proteins, it was plausible that TgATG9 and TgPROP1 could function analogously with a putative TgATG2 at this step of the pathway. The identification of a candidate TgATG2 protein in *T. gondii* was based on analysis of domain similarities to other ATG2 proteins and predictive structural modeling using AlphaFold. The putative TgATG2 contained a lipid-transferase-like domain that was predicted to fold into a rod-like tubular groove lined with hydrophobic residues, in which phospholipids could potentially be shuttled. Surprisingly, the putative TgATG2 appears to be five times larger than human and yeast ATG2/Atg2 proteins and larger also than other lipid-transferases, such as those of the Vps13 family, for which several orthologues can be found in the *T. gondii* genome. Vps13 shares high sequence similarity with Atg2 and has been shown to mediate lipid transfer at various organelle contact sites (77–79). In contrast, there is only a single homolog of Atg9 in *T. gondii*. Exploring the role of Vps13-like proteins in *T. gondii* and how they may relate to TgATG2 and autophagy should be the focus of future studies. In this work, we show that TgATG2 interacts with both TgATG9 and TgPROP1, further supporting the candidacy of TgATG2 as a bona fide autophagy protein. While our work has identified interactions among potential key proteins of a putative membrane expansion complex in *T. gondii* autophagy, future studies are necessary to define the mechanism of action of TgATG9–TgATG2–TgPROP1 in the expansion step of autophagy. Furthermore, mapping the exact sites of interaction will reveal key insights into potential similarities or differences in how the expansion complex is formed in *T. gondii* compared to other model systems (15, 80–82). Additional studies are also needed to determine if this complex is positioned between donor membranes, such as the endoplasmic reticulum, and the developing autophagic membrane. While our immunofluorescence assays provide a snapshot of the subcellular localization of TgATG2, TgATG9 and TgPROP1 within tissue culture derived bradyzoites, higher resolution microscopy techniques such as lattice light sheet microscopy (83–85), may be necessary to further elucidate the precise subcellular dynamics of these proteins.

Although we did not investigate whether this putative TgATG2 has lipid-transferase activity, we generated two independent TgATG2 knockouts (TgATG2 KO and TgATG2′ KO) and demonstrated that TgATG2 is required for extracellular tachyzoite fitness and cyst persistence, consistent with observations for TgATG9 (41, 42). Because tachyzoites survive only briefly outside host cells, even modest losses in extracellular viability can have major downstream effects; accordingly, TgATG2-deficient parasites exhibited reduced extracellular survival, which was accompanied by impaired invasion and fewer plaques. While the mechanistic basis of this phenotype was not directly examined here, we propose that TgATG2 loss diminishes autophagic flux, thereby impairing the ability of tachyzoites to tolerate the transient stresses associated with extracellular exposure before establishing a new intracellular niche.

We further confirmed the importance of TgATG2 for bradyzoite autophagy by assessing CytoID-positive autolysosome accumulation upon LHVS treatment in both TgATG2 knockout strains. Ablation of TgATG2 caused a marked reduction in the accumulation of CytoID-positive autophagolysosomes. Although our previous studies have relied heavily on CytoID staining to assess bradyzoite autophagy, this approach does not quantify flux directly and instead requires LHVS to promote the buildup of autophagic cargo in compartments detected by the dye. We addressed these limitations by developing a dual-fluorescence TgATG8 reporter that directly measured autophagic flux in *T. gondii* bradyzoites. The reporter was validated using TgATG9 KO parasites, which are known to have a strong defect in autophagy, and which resulted in a significant increase in the GFP:tdTomato ratio indicative of decreased autophagic flux. Furthermore, western blot analysis showed that free GFP was absent, suggesting that autophagy was dramatically reduced or inactive. We further tested the reporter by deleting the whole TgATG2 gene or partially deleting it, following the same approach as for the previous TgATG2 KO strains.

TgATG2 ablation caused an increase in the GFP:tdTomato ratio in both KO strains, indicative of decreased flux. Residual autophagic activity in the absence of TgATG2 was quantified by measuring free GFP levels, which were approximately 30–35% of those observed in the WT. Therefore, our reporter provides an alternative to the indirect interpretations of autophagic flux status within bradyzoites offered by the CytoID assay. Validation with other chemical modulators of autophagy widely used in other systems was not feasible in *T. gondii*, as many targets are absent, such as the target of rapamycin (TOR), or are known to have off-target effects (such as chloroquine or bafilomycin A1) (86–91). Nonetheless, our new reporter could be used in future high-throughput chemical or genetic screens to potentially identify novel inhibitors of *T. gondii* autophagy or new candidate TgATGs, respectively.

Finally, we confirmed the necessity of TgATG2 and autophagy for bradyzoite survival using a mouse model of chronic *T. gondii* infection. Mice chronically infected with TgATG2 KOs exhibited a significant reduction in cyst burdens in the brain and decreased bradyzoite viability of *ex vivo* cysts, as observed in in vitro bradyzoites. Interestingly, the cyst burden reduction of TgATG2 KOs was not as severe as the reduction seen for TgATG9 KO (41).We still detected some cysts in brains infected with TgATG2 KOs, whereas TgATG9 KO-infected brains had barely any detectable cysts. The *in vivo* results were in line with the reduction in autophagosomes observed in the *in vitro* experiments, in which some CytoID staining was still observed in the cysts of the two TgATG2 KOs, although at very low intensity. A potential explanation for these observed phenotypes is redundancy among lipid-transferase proteins such as Vps13, for which several orthologues can be found in the *T. gondii* genome.

Overall, our work extends the limited knowledge of known TgATGs through identification of TgATG2, which may form the basis of a membrane expansion complex critical for *T. gondii* bradyzoite autophagy.

## MATERIAL AND METHODS

### Cell culture, parasite maintenance, and bradyzoite differentiation

*Toxoplasma gondii* tachyzoites were propagated in vitro by serial passage on human foreskin fibroblast (HFF) monolayers (58, 92). HFFs were maintained at 37°C in a humidified 5% CO₂ incubator and cultured in high-glucose Dulbecco’s Modified Eagle Medium (DMEM; EuroClone) supplemented with 10% heat-inactivated fetal bovine serum (Thermo Fisher Scientific), 2 mM L-glutamine (Euroclone), and 50 µg/mL penicillin-streptomycin (Euroclone), hereafter designated as D10 medium (93). To induce in vitro bradyzoite differentiation, tachyzoites were mechanically harvested from infected HFFs via scraping and passage through 26-gauge needles and subsequently inoculated onto fresh HFF monolayers. Following a 24-h invasion period at 37°C under 5% CO₂ in D10 medium, tachyzoite-to-bradyzoite conversion was stimulated by replacing the culture medium with alkaline differentiation medium. This medium consisted of NaHCO₃-free RPMI-1640 (Cytiva, SH30011.02) supplemented with 3% heat-inactivated fetal bovine serum, 50 mM HEPES (Sigma-Aldrich, H3375), and 50 µg/mL penicillin-streptomycin, adjusted to pH 8.2–8.3 with NaOH. Differentiation cultures were maintained at 37°C under ambient CO₂ conditions, with daily medium replacement to sustain the alkaline pH required to drive encystation (73, 94).

### Generation of transgenic parasite lines

All transgenic *Toxoplasma gondii* strains were generated in either ME49/DD-BFD1 or ME49Δku8Δhxgprt parental background via CRISPR/Cas9-mediated homologous recombination. Briefly, starting with the DD-BFD1 expressing ME49 *T. gondii* strain (ME49/DD-BFD1) (51), TgATG2 was endogenously tagged at the C terminus to generate ME49/DD-BFD1/TgATG2-smHA parasites. A guide RNA targeting the C terminus of TgATG2 near the stop codon was generated, and the homology-directed repair template was produced by PCR amplification of spaghetti-monster-HA-CAT. ME49/DD-BFD1 parasites were co-transfected with 100 µg of guide RNA and 5 µg of repair template, as described previously (92, 94, 95). At 24 h post-transfection, positively transfected parasites were selected with 20 µM chloramphenicol before clonal isolation by limiting dilution. Individual parasite clones were validated by PCR amplification to confirm the presence of the smHA tag (96). Subsequently, TgATG9 was endogenously tagged at the C terminus using a similar strategy to generate ME49/DD-BFD1/TgATG2-smHA/TgATG9-smMyc parasites. Positively transfected parasites were selected with 5 µg/mL phleomycin (Invitrogen, NC9198593) before clonal isolation by limiting dilution, and individual clones were validated by PCR amplification to confirm the presence of the smMyc tag. The ME49/DD-BFD1/TgPROP1-smHA/TgATG2-smMyc strain was generated using the same strategy used for ME49/DD-BFD1/TgATG2-smHA/TgATG9-smMyc, and individual parasite clones were validated by PCR amplification to confirm the presence of the smMyc tag. Generation of the TgATG2-smHA-tagged strain in the ME49Δku80Δhxgprt background followed the same strategy described above. To generate the MΔTgATG2 line, the entire genomic sequence of TgATG2 was replaced with the chloramphenicol acetyltransferase (CAT) cassette using two guide RNAs targeting the N and C termini near the start and stop codons, respectively (Fig. S1). The repair template was amplified using the oligonucleotides listed in Table S1. Conversely, the independent MΔTgATG2’ line was generated by replacing the N-terminal half of TgATG2 with the hypoxanthine-xanthine-guanine phosphoribosyltransferase (HXGPRT) gene, using the same N-terminal guide RNA in combination with a novel guide RNA targeting the middle of the locus (Fig. S1). The repair template was amplified using the oligonucleotides listed in Table S1. For both knockouts, parasites were co-transfected with 100 µg of each guide RNA and 5 µg of the respective repair template. Selection was maintained for 5 days with either 20 µM chloramphenicol or a combination of 25 µg/mL mycophenolic acid and 50 µg/mL xanthine (Sigma-Aldrich), before single-clone isolation. Guide RNAs and oligonucleotides used for genetic manipulations are summarized in Table S1. Clonal parasite lines were isolated by limiting dilution and validated by PCR using clone-specific primers.

For clarity and concise nomenclature throughout this manuscript, the engineered strains are designated as follows: ME49Δku80Δhxgprt as ME49; ME49Δku80ΔhxgprtΔTgATG2 as MΔTgATG2; ME49Δku80ΔhxgprtΔTgATG2’ as MΔTgATG2’; the tdTomato-2A-GFP-TgATG8 autophagic flux reporter background as M/Sn; and the reporter-containing TgATG9, TgATG2, and TgATG2’ knockout lines as M/SnΔTgATG9, M/SnΔTgATG2, and M/SnΔTgATG2’, respectively.

### Co-immunoprecipitation

Double-tagged ME49/DD-BFD1/TgATG2-smHA/TgATG9-smMyc or ME49/DD-BFD1/TgPROP1-smHA/TgATG2-smMyc parasites and single-tagged and untagged controls were differentiated in 150 mm tissue culture dishes (Corning, CLS430599) for 7 days in D10 medium supplemented with 3 μM Shield-1. To generate enough bradyzoite material for co-immunoprecipitation, at least six 150 mm dishes were needed per parasite strain. On day 7, bradyzoites were harvested via scraping, syringing (20-and 25-gauge needles), and resuspended in 1 mL of Pierce IP lysis buffer (Thermo Scientific, 87787) supplemented with complete Mini Protease Inhibitors cocktail (Roche, 11836153001). Lysates were centrifuged at 4°C for 10 min at 14,000 x g. Prior to immunoprecipitation, 10% of each lysate was saved as the “input”. For co-immunoprecipitation using smHA-tagged proteins as the bait (TgATG2-smHA or TgPROP1-smHA), equal volumes (∼500 μL) of lysates were incubated with 5 μL of rat anti-HA (Roche, 11867423001) or without antibody as a control for 1 hour at 4°C, nutating. After 1 hour, 50 μL of a 50% slurry of Protein A Sepharose 4B beads (Thermo Scientific, 101041) washed with Pierce IP lysis buffer was added to each lysate for 1 hour, rocking at room temperature. Beads were pelleted for 2 min at 4000 rpm followed by 3 washes with 1 mL IP lysis buffer prior to elution. Proteins were eluted with 90 μL of a low pH glycine-HCl elution buffer (0.1 M, pH 2.3) and incubated with frequent agitation for 10 min at room temperature. Beads were pelleted for 2 min at 4000 rpm, then supernatants neutralized with 10 μL of 1M Tris pH 8.0-9.0. Proteins were precipitated using methanol-chloroform precipitation prior to western or dot blotting. For western blot after co-immunoprecipitation, inputs, washes, and eluted proteins were supplemented with 5X SDS-PAGE sample buffer and 10% β-mercaptoethanol, resulting in a final concentration of 1X SDS-PAGE buffer and 2% β-mercaptoethanol. For dot blot after co-immunoprecipitation, samples were added to a 96-well Bio-Dot Apparatus (Biorad, 1706545) containing a 0.45-μm nitrocellulose membrane that had been prepared for dot blotting procedures according to the manufacturer’s manual. Samples were allowed to filter through the membrane by gravity flow for 1.5 hours at room temperature. After filtering through samples by gravity flow, wells were washed 3 times with PBS-T prior to removing the nitrocellulose membrane from the Bio-Dot Apparatus. Membranes were blocked for 30 min at room temperature in phosphate-buffered saline containing 0.05% Tween-20 and 5% milk and then processed as described for western blotting.

### Western blot analysis

To validate TgATG2 knockout, parasites were differentiated into bradyzoites for 7 days under alkaline stress with daily medium replacement, followed by a 3-day treatment with either 1 μM morpholinurea-leucine-homophenylalanine-phenyl-vinyl-sulfone (LHVS) or DMSO vehicle control when indicated. Bradyzoites were harvested by scraping, syringe passage (26-gauge needles), and pepsin digestion (0.026% pepsin in 170 mM NaCl and 60 mM HCl) for 30 min at 37°C. Parasites were then enumerated and lysed in RIPA buffer (Thermo Fisher Scientific) supplemented with protease inhibitors (Roche) for 30 min on ice. Lysates were cleared by centrifugation (16,000 × g, 30 min, 4°C), denatured in 5× Laemmli buffer, and boiled at 98°C for 5 min. Equivalent lysates (∼ 3.6 × 10⁶ bradyzoites per lane) were resolved on 12% SDS-PAGE gels and transferred onto PVDF membranes using a Trans-Blot Turbo semi-dry transfer system (Bio-Rad). Membranes were blocked in 5% nonfat dry milk in Tris-buffered saline containing 0.1% Tween 20 (TBST) for 1 h at room temperature and incubated overnight at 4°C with primary antibodies diluted in 2.5% milk/TBST. Mouse anti-EGFP (Thermo Fisher Scientific) was used 1:1,000; rabbit anti-BAG1 1:5,000; rabbit anti-Myc (Cell Signaling Technology, 2278) 1:1000; rabbit anti-TgActin (Sibley lab, Washington University in St. Louis) 1:20,000; rat anti-HA (Roche, 11867423001) 1:2500; mouse anti-HA (Thermo Fisher Scientific) 1:1,000. After primary antibody incubation, membranes were washed 3 times with PBS-T before incubation with HRP-conjugated secondary antibodies (Jackson ImmunoResearch Laboratories, 115-035-146; 1:5000) for 1 h at room temperature. Immunoreactive bands were developed using SuperSignal West Pico PLUS Chemiluminescent Substrate (Thermo Fisher Scientific) and captured on an iBright 1500 system (Thermo Fisher Scientific). Band intensities were quantified and normalized to BAG1 or TgActin loading controls using iBright Analysis Software.

### Quantitative Western blot analysis of GFP processing

Densitometric quantification of immunoreactive bands corresponding to GFP-TgATG8 and free GFP was performed using iBright Analysis Software (Thermo Fisher Scientific) by recording the background-corrected integrated intensity values (“Rolling bg. corr. vol.”) for each target. To control for variations in protein loading and bradyzoite differentiation efficiency, both GFP-TgATG8 and free GFP signals were normalized to the corresponding BAG1 signal from the same lane, which served as the internal loading control.

For each experimental group—comprising Me49/Sn, Me49/SnΔATG9, Me49/SnΔATG2 and Me49/SnΔATG2’ parasites cultured under bradyzoite-inducing conditions in the presence of either DMSO or LHVS—the proportion of processed GFP was calculated using the following equation:

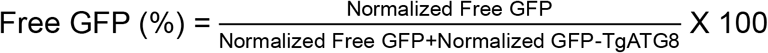

To isolate and quantify autophagy-dependent GFP processing, the baseline percentage of free GFP measured in DMSO-treated controls was subtracted from the values obtained under LHVS-treated conditions:

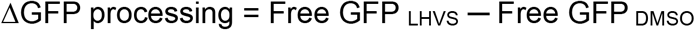

The resulting ΔGFP value reflects the magnitude of LHVS-sensitive autophagic flux. To facilitate direct comparison among the different mutant strains, autophagic activity was expressed relative to the Me49/Sn control, which was defined as 100%:

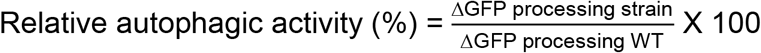

### Construction of the autophagic flux reporter and derived lines

The plasmid used to express the autophagic flux reporter was generated from the pTub-tdTomato-TgATG8-CAT plasmid [9]. The BAG1 promoter was amplified from *T. gondii* ME49 genomic DNA using primers P20/P21 and cloned into the SpeI/XmaI restriction sites of the pTub-tdTomato-TgATG8-CAT plasmid, upstream of the ATG8 coding sequence. From the resulting plasmid, a backbone containing pBAG1 and TgATG8 was amplified using primers P28/P29. A custom plasmid containing the tdTomato-2A-GFP cassette was synthesized by GeneUniversal and the cassette was amplified using primers P30/P31. The backbone and insert fragments were joined using Gibson Assembly (New England Biolabs, E5510S) prior to bacterial transformation. The CAT resistance was then replaced with the Tub/DHFR resistance cassette using the restriction enzymes HindIII/SpeI. The plasmid was then linearized with BcI and integrated into the BAG1 genomic locus of the parental strain via single-crossover homologous recombination (Fig. S2). Transfected parasites were selected using 1 μM pyrimethamine and cloned. This M/Sn reporter background was subsequently utilized to engineer independent mutant lines, including M/SnΔTgATG9, M/SnΔTgATG2 and M/SnΔTgATG2’, applying the CRISPR/Cas9 strategies as described above (Fig. S2). Genetic complementation lines were established in the respective mutant backgrounds by integrating full-length coding sequences under either endogenous or heterologous promoters via single-crossover homologous recombination (97).

### Immunofluorescence microscopy

For immunofluorescence assays (IFA), samples were fixed in 4% methanol-free paraformaldehyde (PFA) in PBS for 20 min, permeabilized with 0.2% Triton X-100 for 20 min, and blocked with 10% fetal bovine serum (FBS) in PBS for 1 h. Primary and secondary antibodies were diluted in 2% FBS/PBS, and all incubations were performed for 1 h at room temperature. Primary antibodies included rat anti-HA (Roche, 11867423001; 1:250) and mouse anti-TgSERCA (1:1,000). Specific signal was visualized using goat anti-rat Alexa Fluor 594 and goat anti-mouse Alexa Fluor 488 secondary antibodies (Invitrogen; both at 1:1,000) (97).

### TgATG8 autophagic flux reporter assay and image quantification

Parasitic autophagic flux was assessed using *T. gondii* lines expressing a tdTomato-2A-GFP-TgATG8 reporter integrated at the BAG1 locus. Tachyzoites harvested from infected HFF monolayers were inoculated onto glass coverslips at a density of ∼10^5^ parasites per well. At 24 h post-infection, tachyzoite-to-bradyzoite differentiation was induced under alkaline conditions for 7 days, with daily medium replacement as described above. Following differentiation, coverslips were washed three times with HBSS supplemented with 1 mM CaCl_2 a_nd 1 mM MgCl_2,_ fixed in 4% methanol-free paraformaldehyde (PFA), and mounted in Mowiol. Fluorescence images were captured on a Zeiss Axio Observer Z1 inverted microscope using a 60× oil immersion objective under identical acquisition settings across all experimental groups. For each parasite strain, a minimum of 500 total cysts were evaluated across three independent biological replicates, each comprising three technical replicates. Image processing and morphometric quantifications were performed using Zeiss ZEN 3.5 (Blue Edition) software. Cysts were manually outlined using the contour tool to extract single cyst mean fluorescence intensities for the GFP (FITC) and tdTomato (TRITC) channels. Autophagic flux was quantified as the FITC/TRITC fluorescence intensity ratio. For each biological replicate, mean ratios were computed and normalized to untreated wild-type (WT) controls prior to statistical evaluation.

### CytoID staining and quantification of autolysosomes

*Toxoplasma gondii* tachyzoites were seeded onto 22 × 22 mm (No. 1.5) glass coverslips in six-well plates and induced to differentiate into bradyzoites under alkaline-stress conditions for 7 days. Where indicated, bradyzoites were treated with either 1 μM LHVS or an equivalent volume of DMSO vehicle control for an additional 3 days. For genetic ablation experiments, MΔTgATG2 and MΔTgATG2′ control parasites were subjected to identical treatment regimens. Autolysosomes were labeled in live parasites using the CytoID Autophagy Detection Kit 2.0 (Enzo Life Sciences, ENZ-KIT175) according to the manufacturer’s instructions (24). Following a 1-h incubation with the CytoID reagent, samples were fixed with 4% paraformaldehyde and mounted using ProLong Gold Antifade Mountant (Thermo Fisher Scientific) or Mowiol (Sigma-Aldrich). Fluorescence images were captured on a Nikon Eclipse TE2000-U microscope using identical acquisition settings across all experimental groups, and image analysis was performed using ZEN 3.7 Blue Edition software.

### PMA-qPCR viability assays

To quantify parasite viability based on membrane integrity, a propidium monoazide (PMA)-based qPCR assay was adapted from established protocols (58). PMAxx (Biotium) selectively penetrates membrane-compromised cells and covalently cross-links to genomic DNA upon photoactivation at 470 nm, thereby inhibiting PCR amplification. To minimize processing variability, qPCR was performed directly on cell lysates without prior DNA purification using DNARelease Additive (Phire Tissue Direct PCR Master Kit; Thermo Fisher Scientific).

For extracellular tachyzoites, freshly egressed parasites were purified from host debris using 3-μm filters, washed thrice, and resuspended in Hank’s Balanced Salt Solution (HBSS) at 37°C. Aliquots collected at designated intervals (0, 30, 60, and 180 min) were incubated with or without 30 μM PMAxx for 15 min in the dark with gentle agitation, followed by 15 min of blue-light exposure using a PMA-Lite LED Photolysis Device (Biotium). For bradyzoites, infected HFF monolayers in 96-well plates (1 × 10² tachyzoites/well) were cultured in alkaline differentiation medium for 1 or 2 weeks. Bradyzoites were released by pepsin digestion (0.026% pepsin in 170 mM NaCl and 60 mM HCl) for 1 h at 37°C, neutralized with an equal volume of 188 mM Na₂CO₃, and treated on ice with either 30 μM PMAxx or a ddH₂O control under identical photolysis conditions. Following photoactivation, lysates were prepared by adding 1 μL of DNARelease Additive, followed by sequential incubations at 22°C for 4 min and 98°C for 2 min. Real-time qPCR was performed using 1 μL of lysate, PowerTrack SYBR Green Master Mix (Thermo Fisher Scientific), and 0.3 μM of multilocus primers (T9-T11 see **Table S1**) under the following cycling conditions: 95°C for 15 s and 60°C for 30 s for 40 cycles. Parasite viability was calculated using the following equations:

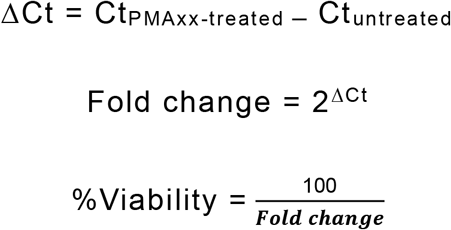

### Plaque assay

Extracellular *Toxoplasma gondii* tachyzoites were mechanically harvested from infected HFF monolayers via scraping and syringe passage (25-gauge needle), followed by purification from host cell debris. Parasites were enumerated using a hemocytometer and inoculated onto confluent HFF monolayers in six-well plates at a density of 300 parasites per well, utilizing triplicate or quadruplicate technical replicates. Following infection, plates were cultured in complete D10 medium supplemented with 10 mM HEPES and maintained undisturbed at 37°C under 5% CO₂ for 14 days to permit plaque development. Monolayers were subsequently fixed and stained with a crystal violet solution, gently rinsed with water, and air-dried. Lysis plaques, identified as clear zones within the stained monolayer, were quantified using the Viral Plaque macro in ImageJ. All assays were executed across at least three independent biological replicates.

### Invasion assay

Host cell invasion assays were performed as previously described utilizing a red–green differential immunofluorescence strategy (98). Briefly, extracellular (attached) tachyzoites were labeled in non-permeabilized samples using an anti-TgSAG1 antibody. Following cell permeabilization with 0.2% Triton X-100, the total parasite population was visualized by staining with an anti-GAP45 antibody. Samples were analyzed via fluorescence microscopy, evaluating 300 random fields per well under 100× magnification. Data are representative of three independent biological replicates, each performed in triplicate.

### Mouse infection and brain cyst analysis

Eight-week-old female CBA/J mice (Jackson Laboratory, 000656) were randomly assigned to experimental groups and infected intraperitoneally with 500 tachyzoites of the indicated *Toxoplasma gondii* ME49-derived strains (ME49ΔTgATG2, ME49ΔTgATG2′). All animal procedures were performed in an AAALAC-accredited facility in accordance with the United States Public Health Service guidelines and were approved by the Institutional Animal Care and Use Committee of the University of Michigan (PRO00010428; A3114-01). At 5 weeks post-infection, mice were humanely euthanized and brains were harvested. Brain tissue was homogenized in 1 mL of ice-cold PBS by mechanical disruption using scissors followed by serial passage through 20–21G syringe needles. Samples were blinded prior to quantification. Tissue cysts were enumerated by light microscopy from three 10 μL aliquots per brain, and the total cyst burden per brain was calculated by extrapolation to the total homogenate volume.

### *Ex vivo* bradyzoite viability assay

*Ex vivo* bradyzoite viability was assessed from the brains of chronically infected mice. Brain tissue was homogenized in sterile Hanks’ Balanced Salt Solution (HBSS; Gibco, 14175103), and bradyzoites were released by pepsin digestion (0.026% pepsin in 170 mM NaCl and 60 mM HCl; Sigma-Aldrich, P6887) for 30 min at 37°C (73). Parasite viability was determined by combining plaque formation assays with qPCR-based normalization of the initial parasite input. Pepsin-released bradyzoites were inoculated in equal volumes in triplicate onto confluent HFF monolayers in six-well plates containing D10 medium and incubated undisturbed at 37°C under 5% CO₂ for 14 days. Plaques were fixed and stained with a 0.2% crystal violet solution in 70% ethanol for 20 min at room temperature and subsequently quantified via light microscopy. In parallel, genomic DNA was extracted from aliquots of the pepsin-treated samples using the DNeasy Blood and Tissue Kit (Qiagen, 69506). Quantitative PCR was performed using SsoAdvanced Universal SYBR Green Supermix (Bio-Rad, 172-5271) with primers targeting the 529-bp repetitive element of *T. gondii* (58). Reactions were executed on a CFX96 Touch Real-Time PCR Detection System (Bio-Rad) under the following conditions: initial denaturation at 98°C for 3 min, followed by 40 cycles of 98°C for 15 s, 58.5°C for 30 s, and 72°C for 30 s. A standard curve ranging from 6.4 to 2,000 parasite genome equivalents was utilized to calculate absolute genome input. Final plaque numbers were normalized to these genome equivalents to determine relative bradyzoite viability.

### Statistical analysis

Statistical analyses were performed using GraphPad Prism, with outliers identified and excluded via the ROUT method (**Q** = 0.1%). Dataset normality and homogeneity of variance were systematically evaluated using the D’Agostino–Pearson omnibus test and appropriate variance criteria prior to hypothesis testing. For normally distributed data exhibiting equal variances, treatment groups were compared using Student’s *t*-test or one-way/two-way analysis of variance (ANOVA), followed by appropriate post-hoc tests. For non-normally distributed datasets, nonparametric alternatives (Mann–Whitney U or Kruskal–Wallis tests) were applied. Sample sizes were determined based on technical assay capacity and historical experimental variance, with specific statistical tests and sample numbers (**n**) detailed in the respective figure legends. No formal power analysis was conducted.

## ACKNOWLEDGMENTS

This study was supported by National Institutes of Health grants R01AI120627 (to V.B.C. and M.D.C.). We thank My-Hang Huynh for assisting with some experiments and Fengrong Wang for helping to design the autophagic flux reporter plasmid.

## AUTHOR CONTRIBUTIONS

VBC and MDC conceived the study. SM and PT performed most of the experiments and analyzed the data. FP generated the TgATG2 KO strain. TLS made mouse infections. SM, PT, MDC and VBC made the figures. MDC and SM wrote the paper and SM, PT and VBC revised the paper.

## SUPPLEMENTARY LEGENDS

**Figure S1.** Genetic ablation of TgATG2 was achieved with CRISPR-Cas9 and homology-directed insertion of either the hypoxanthine-xanthine-guanine phosphoribosyl transferase (HXGPRT) gene or Chloramphenicol Acetyl Transferase (CAT) into the TgATG2 locus (KO). Two independent TgATG2 KO strains were generated in Me49Δku80 by removing the first part of (MΔTgATG’, left panel) or the entire (MΔTgATG2, right panel) TgATG2 gene, respectively. Bottom panels show PCR validation for the insertion of the HXGPRT or CAT gene in the TgATG2 gene in the KO clone isolated.

**Figure S2.** Strategy used to generate the M/Sn strain and MΔTgATG9, MΔTgATG2 or MΔTgATG2’. The Autophagy flux sensor pBAG1-tdTomato-2A-GFP-TgATG8 plasmid was first linearized using the restriction enzyme SapI and then integrated by single crossing-over at the Bag1 promoter, resulting in the duplication of the Bag1 promoter which now expresses both tdTomato-2A-GFP-TgATG8 and BAG1. This basal strain was used to generate MΔTgATG9, MΔTgATG2 or MΔTgATG2’ using the same strategy of the previous strains. On the top right PCR validation for the insertion of the pBAG1-tdTomato-2A-GFP-TgATG8 plasmid into the Bag1 locus. Bottom panels show PCR validation for the insertion of the selection cassettes in the TgATG9 or TgATG2 gene in the KO clone isolated.

**Table S1. gRNAs and primers used in this study.**

All gRNA and primer sequences are shown in the 5′ to 3′ orientation. Primers are listed according to their names and corresponding PCR targets. The specific purpose of each primer is indicated in the schematic representations of the strategies used to generate and validate knockout and endogenously tagged *Toxoplasma gondii* strains.

## REFERENCES

1. Reggiori F, Klionsky DJ. 2002. Autophagy in the eukaryotic cell. Eukaryot Cell 1:11–21.

2. Mizushima N, Komatsu M. 2011. Autophagy: renovation of cells and tissues. Cell 147:728–741.

3. Lei Y, Huang Y, Wen X, Yin Z, Zhang Z, Klionsky DJ. 2022. How Cells Deal with the Fluctuating Environment: Autophagy Regulation under Stress in Yeast and Mammalian Systems. Antioxidants (Basel) 11:304.

4. Matoba K, Noda NN. 2021. Structural catalog of core Atg proteins opens new era of autophagy research. J Biochem 169:517–525.

5. Feng Y, He D, Yao Z, Klionsky DJ. 2014. The machinery of macroautophagy. Cell Res 24:24–41.

6. Hurley JH. 2026. The Human Autophagy Core Complexes. Annu Rev Biochem 95:507–524.

7. Fujioka Y, N. Noda N. 2025. Mechanisms of autophagosome formation. Proc Jpn Acad Ser B Phys Biol Sci 101:32–40.

8. Sawa-Makarska J, Baumann V, Coudevylle N, von Bülow S, Nogellova V, Abert C, Schuschnig M, Graef M, Hummer G, Martens S. 2020. Reconstitution of autophagosome nucleation defines Atg9 vesicles as seeds for membrane formation. Science 369:eaaz7714.

9. Noda NN. 2021. Atg2 and Atg9: Intermembrane and interleaflet lipid transporters driving autophagy. Biochim Biophys Acta Mol Cell Biol Lipids 1866:158956.

10. Valverde DP, Yu S, Boggavarapu V, Kumar N, Lees JA, Walz T, Reinisch KM, Melia TJ. 2019. ATG2 transports lipids to promote autophagosome biogenesis. J Cell Biol 218:1787–1798.

11. Swan LE. 2025. VPS13 and bridge-like lipid transporters, mechanisms, and mysteries. Front Neurosci 19:1534061.

12. Neuman SD, Levine TP, Bashirullah A. 2022. A novel superfamily of bridge-like lipid transfer proteins. Trends Cell Biol 32:962–974.

13. Kotani T, Kirisako H, Koizumi M, Ohsumi Y, Nakatogawa H. 2018. The Atg2-Atg18 complex tethers pre-autophagosomal membranes to the endoplasmic reticulum for autophagosome formation. Proc Natl Acad Sci U S A 115:10363–10368.

14. Otomo T, Maeda S. 2019. ATG2A transfers lipids between membranes in vitro. Autophagy 15:2031–2032.

15. Gómez-Sánchez R, Rose J, Guimarães R, Mari M, Papinski D, Rieter E, Geerts WJ, Hardenberg R, Kraft C, Ungermann C, Reggiori F. 2018. Atg9 establishes Atg2-dependent contact sites between the endoplasmic reticulum and phagophores. J Cell Biol 217:2743–2763.

16. Osawa T, Matoba K, Noda NN. 2022. Lipid Transport from Endoplasmic Reticulum to Autophagic Membranes. Cold Spring Harb Perspect Biol 14:a041254.

17. A model for a partnership of lipid transfer proteins and scramblases in membrane expansion and organelle biogenesis | PNAS. https://www.pnas.org/doi/10.1073/pnas.2101562118. Retrieved 6 July 2026.

18. Romano PS, Akematsu T, Besteiro S, Bindschedler A, Carruthers VB, Chahine Z, Coppens I, Descoteaux A, Alberto Duque TL, He CY, Heussler V, Le Roch KG, Li F-J, de Menezes JPB, Menna-Barreto RFS, Mottram JC, Schmuckli-Maurer J, Turk B, Tavares Veras PS, Salassa BN, Vanrell MC. 2023. Autophagy in protists and their hosts: When, how and why? Autophagy Rep 2:2149211.

19. Fu J, Zhao L, Pang Y, Chen H, Yamamoto H, Chen Y, Li Z, Mizushima N, Jia H. 2023. Apicoplast biogenesis mediated by ATG8 requires the ATG12-ATG5-ATG16L and SNAP29 complexes in Toxoplasma gondii. Autophagy 19:1258–1276.

20. Cheng L, Tian Y, Wang Y, Wang T, Yao Y, Yu H, Zheng X, Wu M, Zhao W, Hua Q, Hu X, Tan F. 2022. Toxoplasma TgAtg8-TgAtg3 Interaction Primarily Contributes to Apicoplast Inheritance and Parasite Growth in Tachyzoite. Microbiol Spectr 10:e0149521.

21. Besteiro S. 2017. Autophagy in apicomplexan parasites. Curr Opin Microbiol 40:14–20.

22. Bansal P, Tripathi A, Thakur V, Mohmmed A, Sharma P. 2017. Autophagy-Related Protein ATG18 Regulates Apicoplast Biogenesis in Apicomplexan Parasites. mBio 8:e01468–17.

23. Nguyen HM, Berry L, Sullivan WJ, Besteiro S. 2017. Autophagy participates in the unfolded protein response in Toxoplasma gondii. FEMS Microbiol Lett 364:fnx153.

24. Di Cristina M, Dou Z, Lunghi M, Kannan G, Huynh M-H, McGovern OL, Schultz TL, Schultz AJ, Miller AJ, Hayes BM, van der Linden W, Emiliani C, Bogyo M, Besteiro S, Coppens I, Carruthers VB. 2017. Toxoplasma depends on lysosomal consumption of autophagosomes for persistent infection. Nat Microbiol 2:17096.

25. Latré de Laté P, Pineda M, Harnett M, Harnett W, Besteiro S, Langsley G. 2017. Apicomplexan autophagy and modulation of autophagy in parasite-infected host cells. Biomedical Journal 40:23–30.

26. Nguyen HM, El Hajj H, El Hajj R, Tawil N, Berry L, Lebrun M, Bordat Y, Besteiro S. 2017. Toxoplasma gondii autophagy-related protein ATG9 is crucial for the survival of parasites in their host. Cell Microbiol 19.

27. Lévêque MF, Berry L, Cipriano MJ, Nguyen H-M, Striepen B, Besteiro S. 2015. Autophagy-Related Protein ATG8 Has a Noncanonical Function for Apicoplast Inheritance in Toxoplasma gondii. mBio 6:e01446–01415.

28. Sinai AP, Roepe PD. 2012. Autophagy in Apicomplexa: a life sustaining death mechanism? Trends in Parasitology 28:358–364.

29. Besteiro S. 2012. Which roles for autophagy in Toxoplasma gondii and related apicomplexan parasites? Mol Biochem Parasitol 184:1–8.

30. Ghosh D, Walton JL, Roepe PD, Sinai AP. 2012. Autophagy is a cell death mechanism in Toxoplasma gondii. Cell Microbiol 14:589–607.

31. Kong-Hap MA, Mouammine A, Daher W, Berry L, Lebrun M, Dubremetz J-F, Besteiro S. 2013. Regulation of ATG8 membrane association by ATG4 in the parasitic protist Toxoplasma gondii. Autophagy 9:1334–1348.

32. Sakamoto H, Nakada-Tsukui K, Besteiro S. 2021. The Autophagy Machinery in Human-Parasitic Protists; Diverse Functions for Universally Conserved Proteins. Cells 10:1258.

33. Wu M, Ying J, Lin X, Xu C, Zheng X, Zheng Y, Fang Z, Yan B, Zhang N, Mou Y, Tan F. 2024. Toxoplasma gondii autophagy-related protein ATG7 maintains apicoplast inheritance by stabilizing and lipidating ATG8. Biochim Biophys Acta Mol Basis Dis 1870:166891.

34. Tomlins AM, Ben-Rached F, Williams RA, Proto WR, Coppens I, Ruch U, Gilberger TW, Coombs GH, Mottram JC, Müller S, Langsley G. 2013. Plasmodium falciparum ATG8 implicated in both autophagy and apicoplast formation. Autophagy 9:1540–1552.

35. Eickel N, Kaiser G, Prado M, Burda P-C, Roelli M, Stanway RR, Heussler VT. 2013. Features of autophagic cell death in Plasmodium liver-stage parasites. Autophagy 9:568–580.

36. Cervantes S, Bunnik EM, Saraf A, Conner CM, Escalante A, Sardiu ME, Ponts N, Prudhomme J, Florens L, Le Roch KG. 2014. The multifunctional autophagy pathway in the human malaria parasite, Plasmodium falciparum. Autophagy 10:80–92.

37. Li Y, Niu Z, Yang J, Yang X, Chen Y, Li Y, Liang X, Zhang J, Fan F, Wu P, Peng C, Shen B. 2023. Rapid metabolic reprogramming mediated by the AMP-activated protein kinase during the lytic cycle of Toxoplasma gondii. Nat Commun 14:422.

38. Yang X, Yang J, Lyu M, Li Y, Liu A, Shen B. 2024. The α subunit of AMP-activated protein kinase is critical for the metabolic success and tachyzoite proliferation of Toxoplasma gondii. Microb Biotechnol 17:e14455.

39. Latré de Laté P, Pineda M, Harnett M, Harnett W, Besteiro S, Langsley G. 2017. Apicomplexan autophagy and modulation of autophagy in parasite-infected host cells. Biomed J 40:23–30.

40. Hain AUP, Bosch J. 2013. Autophagy in Plasmodium, a multifunctional pathway? Comput Struct Biotechnol J 8:e201308002.

41. Smith D, Kannan G, Coppens I, Wang F, Nguyen HM, Cerutti A, Olafsson EB, Rimple PA, Schultz TL, Mercado Soto NM, Di Cristina M, Besteiro S, Carruthers VB. 2021. Toxoplasma TgATG9 is critical for autophagy and long-term persistence in tissue cysts. Elife 10:e59384.

42. Thaprawat P, Zhang Z, Rentchler EC, Wang F, Chalasani S, Giuliano CJ, Lourido S, Di Cristina M, Klionsky DJ, Carruthers VB. 2024. TgATG9 is required for autophagosome biogenesis and maintenance of chronic infection in Toxoplasma gondii. Autophagy Rep 3:2418256.

43. Thaprawat P, Wang F, Chalasani S, Schultz TL, Di Cristina M, Carruthers VB. 2025. Toxoplasma gondii PROP1 is critical for autophagy and parasite viability during chronic infection. mSphere 10:e0082924.

44. Stasic AJ, Moreno SNJ, Carruthers VB, Dou Z. 2022. The Toxoplasma plant-like vacuolar compartment (PLVAC). J Eukaryot Microbiol 69:e12951.

45. Dou Z, McGovern OL, Di Cristina M, Carruthers VB. 2014. Toxoplasma gondii ingests and digests host cytosolic proteins. mBio 5:e01188–01114.

46. Kannan G, Thaprawat P, Schultz TL, Carruthers VB. 2021. Acquisition of Host Cytosolic Protein by Toxoplasma gondii Bradyzoites. mSphere 6:e00934–20.

47. Parussini F, Coppens I, Shah PP, Diamond SL, Carruthers VB. 2010. Cathepsin L occupies a vacuolar compartment and is a protein maturase within the endo/exocytic system of Toxoplasma gondii. Mol Microbiol 76:1340–1357.

48. Besteiro S, Brooks CF, Striepen B, Dubremetz J-F. 2011. Autophagy protein Atg3 is essential for maintaining mitochondrial integrity and for normal intracellular development of Toxoplasma gondii tachyzoites. PLoS Pathog 7:e1002416.

49. Larson ET, Parussini F, Huynh M-H, Giebel JD, Kelley AM, Zhang L, Bogyo M, Merritt EA, Carruthers VB. 2009. Toxoplasma gondii cathepsin L is the primary target of the invasion-inhibitory compound morpholinurea-leucyl-homophenyl-vinyl sulfone phenyl. J Biol Chem 284:26839–26850.

50. Gajria B, Bahl A, Brestelli J, Dommer J, Fischer S, Gao X, Heiges M, Iodice J, Kissinger JC, Mackey AJ, Pinney DF, Roos DS, Stoeckert CJ, Wang H, Brunk BP. 2008. ToxoDB: an integrated Toxoplasma gondii database resource. Nucleic Acids Res 36:D553–556.

51. Waldman BS, Schwarz D, Wadsworth MH, Saeij JP, Shalek AK, Lourido S. 2020. Identification of a Master Regulator of Differentiation in Toxoplasma. Cell 180:359–372.e16.

52. Viswanathan S, Williams ME, Bloss EB, Stasevich TJ, Speer CM, Nern A, Pfeiffer BD, Hooks BM, Li W-P, English BP, Tian T, Henry GL, Macklin JJ, Patel R, Gerfen CR, Zhuang X, Wang Y, Rubin GM, Looger LL. 2015. High-performance probes for light and electron microscopy. Nat Methods 12:568–576.

53. Hortua Triana MA, Márquez-Nogueras KM, Chang L, Stasic AJ, Li C, Spiegel KA, Sharma A, Li Z-H, Moreno SNJ. 2018. Tagging of Weakly Expressed Toxoplasma gondii Calcium-Related Genes with High-Affinity Tags. J Eukaryot Microbiol 65:709–721.

54. Di Cristina M, Carruthers VB. 2018. New and emerging uses of CRISPR/Cas9 to genetically manipulate apicomplexan parasites. Parasitology 145:1119–1126.

55. Kim K, Soldati D, Boothroyd JC. 1993. Gene replacement in Toxoplasma gondii with chloramphenicol acetyltransferase as selectable marker. Science 262:911–914.

56. Klappan AK, Hones S, Mylonas I, Brüning A. 2012. Proteasome inhibition by quercetin triggers macroautophagy and blocks mTOR activity. Histochem Cell Biol 137:25–36.

57. Chan LL-Y, Shen D, Wilkinson AR, Patton W, Lai N, Chan E, Kuksin D, Lin B, Qiu J. 2012. A novel image-based cytometry method for autophagy detection in living cells. Autophagy 8:1371–1382.

58. Piro F, Masci S, Kannan G, Focaia R, Schultz TL, Thaprawat P, Carruthers VB, Di Cristina M. 2024. A Toxoplasma gondii putative amino acid transporter localizes to the plant-like vacuolar compartment and controls parasite extracellular survival and stage differentiation. mSphere 9:e0059723.

59. Klionsky DJ. 2011. For the last time, it is GFP-Atg8, not Atg8-GFP (and the same goes for LC3). Autophagy 7:1093–1094.

60. Torggler R, Papinski D, Kraft C. 2017. Assays to Monitor Autophagy in Saccharomyces cerevisiae. Cells 6:23.

61. Fan S, Dong S, Yao W, Zhang Y, Fan M, Feng S, Wu C, Zhang L, Yi C. 2025. Mec1-mediated Atg9 phosphorylation regulates the PAS recruitment of Atg9 vesicles upon energy stress. Proc Natl Acad Sci U S A 122:e2422582122.

62. Willis SD, Hanley SE, Doyle SJ, Beluch K, Strich R, Cooper KF. 2022. Cyclin C-Cdk8 Kinase Phosphorylation of Rim15 Prevents the Aberrant Activation of Stress Response Genes. Front Cell Dev Biol 10:867257.

63. Shin KD, Lee HN, Chung T. 2014. A revised assay for monitoring autophagic flux in Arabidopsis thaliana reveals involvement of AUTOPHAGY-RELATED9 in autophagy. Mol Cells 37:399–405.

64. Shintani T, Klionsky DJ. 2004. Cargo proteins facilitate the formation of transport vesicles in the cytoplasm to vacuole targeting pathway. J Biol Chem 279:29889–29894.

65. Kang S, Shin KD, Kim JH, Chung T. 2018. Autophagy-related (ATG) 11, ATG9 and the phosphatidylinositol 3-kinase control ATG2-mediated formation of autophagosomes in Arabidopsis. Plant Cell Rep 37:653–664.

66. Kaizuka T, Morishita H, Hama Y, Tsukamoto S, Matsui T, Toyota Y, Kodama A, Ishihara T, Mizushima T, Mizushima N. 2016. An Autophagic Flux Probe that Releases an Internal Control. Mol Cell 64:835–849.

67. Yoshii SR, Mizushima N. 2017. Monitoring and Measuring Autophagy. Int J Mol Sci 18:1865.

68. Nair U, Thumm M, Klionsky DJ, Krick R. 2011. GFP-Atg8 protease protection as a tool to monitor autophagosome biogenesis. Autophagy 7:1546–1550.

69. Kimura S, Noda T, Yoshimori T. 2007. Dissection of the autophagosome maturation process by a novel reporter protein, tandem fluorescent-tagged LC3. Autophagy 3:452–460.

70. Tanida I, Sou Y, Ezaki J, Minematsu-Ikeguchi N, Ueno T, Kominami E. 2004. HsAtg4B/HsApg4B/autophagin-1 cleaves the carboxyl termini of three human Atg8 homologues and delipidates microtubule-associated protein light chain 3-and GABAA receptor-associated protein-phospholipid conjugates. J Biol Chem 279:36268–36276.

71. Liu Z, Chen O, Wall JBJ, Zheng M, Zhou Y, Wang L, Vaseghi HR, Qian L, Liu J. 2017. Systematic comparison of 2A peptides for cloning multi-genes in a polycistronic vector. Sci Rep 7:2193.

72. Kim JH, Lee S-R, Li L-H, Park H-J, Park J-H, Lee KY, Kim M-K, Shin BA, Choi S-Y. 2011. High cleavage efficiency of a 2A peptide derived from porcine teschovirus-1 in human cell lines, zebrafish and mice. PLoS One 6:e18556.

73. Mayoral J, Di Cristina M, Carruthers VB, Weiss LM. 2020. Toxoplasma gondii: Bradyzoite Differentiation In Vitro and In Vivo. Methods Mol Biol 2071:269–282.

74. Nguyen HM, Liu S, Daher W, Tan F, Besteiro S. 2018. Characterisation of two Toxoplasma PROPPINs homologous to Atg18/WIPI suggests they have evolved distinct specialised functions. PLoS One 13:e0195921.

75. Walczak M, Meister TR, Nguyen HM, Zhu Y, Besteiro S, Yeh E. 2023. Structure-Function Relationship for a Divergent Atg8 Protein Required for a Nonautophagic Function in Apicomplexan Parasites. mBio 14:e0364221.

76. Lévêque MF, Nguyen HM, Besteiro S. 2016. Repurposing of conserved autophagy-related protein ATG8 in a divergent eukaryote. Commun Integr Biol 9:e1197447.

77. Lees JA, Reinisch KM. 2020. Inter-organelle lipid transfer: a channel model for Vps13 and chorein-N motif proteins. Curr Opin Cell Biol 65:66–71.

78. McEwan DG, Ryan KM. 2022. ATG2 and VPS13 proteins: molecular highways transporting lipids to drive membrane expansion and organelle communication. FEBS J 289:7113–7127.

79. Dabrowski R, Tulli S, Graef M. 2023. Parallel phospholipid transfer by Vps13 and Atg2 determines autophagosome biogenesis dynamics. J Cell Biol 222:e202211039.

80. van Vliet AR, Chiduza GN, Maslen SL, Pye VE, Joshi D, De Tito S, Jefferies HBJ, Christodoulou E, Roustan C, Punch E, Hervás JH, O’Reilly N, Skehel JM, Cherepanov P, Tooze SA. 2022. ATG9A and ATG2A form a heteromeric complex essential for autophagosome formation. Mol Cell 82:4324–4339.e8.

81. Guardia CM, Tan X-F, Lian T, Rana MS, Zhou W, Christenson ET, Lowry AJ, Faraldo-Gómez JD, Bonifacino JS, Jiang J, Banerjee A. 2020. Structure of Human ATG9A, the Only Transmembrane Protein of the Core Autophagy Machinery. Cell Rep 31:107837.

82. Vargas Duarte P, Reggiori F. 2023. The Organization and Function of the Phagophore-ER Membrane Contact Sites. Contact (Thousand Oaks) 6:25152564231183898.

83. Liffner B, Absalon S. 2024. Expansion microscopy of apicomplexan parasites. Molecular Microbiology 121:619–635.

84. Qiao C, Li Z, Wang Z, Lin Y, Liu C, Zhang S, Liu Y, Feng Y, Yang X, Fu W, Dong X, Guo J, Xu W, Wang X, Jiang T, Meng Q, Wang Q, Dai Q, Li D. 2025. Fast-adaptive super-resolution lattice light-sheet microscopy for rapid, long-term, near-isotropic subcellular imaging. Nat Methods 22:1059–1069.

85. Chen B-C, Legant WR, Wang K, Shao L, Milkie DE, Davidson MW, Janetopoulos C, Wu XS, Hammer JA, Liu Z, English BP, Mimori-Kiyosue Y, Romero DP, Ritter AT, Lippincott-Schwartz J, Fritz-Laylin L, Mullins RD, Mitchell DM, Bembenek JN, Reymann A-C, Böhme R, Grill SW, Wang JT, Seydoux G, Tulu US, Kiehart DP, Betzig E. 2014. Lattice light-sheet microscopy: Imaging molecules to embryos at high spatiotemporal resolution. Science 346:1257998.

86. Gamea GA, Elmehy DA, Salama AM, Soliman NA, Afifi OK, Elkaliny HH, Abo El Gheit RE, El-Ebiary AA, Tahoon DM, Elkholy RA, Shoeib SM, Eleryan MA, Younis SS. 2022. Direct and indirect antiparasitic effects of chloroquine against the virulent RH strain of Toxoplasma gondii: An experimental study. Acta Trop 232:106508.

87. Gao D, Zhang J, Zhao J, Wen H, Pan J, Zhang S, Fang Y, Li X, Cai Y, Wang X, Wang S. 2014. Autophagy activated by Toxoplasma gondii infection in turn facilitates Toxoplasma gondii proliferation. Parasitol Res 113:2053–2058.

88. Moreno SN, Zhong L, Lu HG, Souza WD, Benchimol M. 1998. Vacuolar-type H+-ATPase regulates cytoplasmic pH in Toxoplasma gondii tachyzoites. Biochem J 330 ( Pt 2):853–860.

89. Delorme-Axford E, Guimaraes RS, Reggiori F, Klionsky DJ. 2015. The yeast Saccharomyces cerevisiae: an overview of methods to study autophagy progression. Methods 75:3–12.

90. Roustan V, Jain A, Teige M, Ebersberger I, Weckwerth W. 2016. An evolutionary perspective of AMPK-TOR signaling in the three domains of life. J Exp Bot 67:3897–3907.

91. Tatebe H, Shiozaki K. 2017. Evolutionary Conservation of the Components in the TOR Signaling Pathways. Biomolecules 7:77.

92. Di Cristina M, Ghouze F, Kocken CH, Naitza S, Cellini P, Soldati D, Thomas AW, Crisanti A. 1999. Transformed Toxoplasma gondii tachyzoites expressing the circumsporozoite protein of Plasmodium knowlesi elicit a specific immune response in rhesus monkeys. Infect Immun 67:1677–1682.

93. Galizi R, Spano F, Giubilei MA, Capuccini B, Magini A, Urbanelli L, Ogawa T, Dubey JP, Spaccapelo R, Emiliani C, Di Cristina M. 2013. Evidence of tRNA cleavage in apicomplexan parasites: Half-tRNAs as new potential regulatory molecules of Toxoplasma gondii and Plasmodium berghei. Mol Biochem Parasitol 188:99–108.

94. Lunghi M, Galizi R, Magini A, Carruthers VB, Di Cristina M. 2015. Expression of the glycolytic enzymes enolase and lactate dehydrogenase during the early phase of Toxoplasma differentiation is regulated by an intron retention mechanism. Mol Microbiol 96:1159–1175.

95. Smith D, Lunghi M, Olafsson EB, Hatton O, Di Cristina M, Carruthers VB. 2023. A High-Throughput Amenable Dual Luciferase System for Measuring Toxoplasma gondii Bradyzoite Viability after Drug Treatment. Anal Chem 95:668–676.

96. Piro F, Carruthers VB, Di Cristina M. 2020. PCR Screening of Toxoplasma gondii Single Clones Directly from 96-Well Plates Without DNA Purification. Methods Mol Biol 2071:117–123.

97. Rivera-Cuevas Y, Mayoral J, Di Cristina M, Lawrence A-LE, Olafsson EB, Patel RK, Thornhill D, Waldman BS, Ono A, Sexton JZ, Lourido S, Weiss LM, Carruthers VB. 2021. Toxoplasma gondii exploits the host ESCRT machinery for parasite uptake of host cytosolic proteins. PLoS Pathog 17:e1010138.

98. Possenti A, Di Cristina M, Nicastro C, Lunghi M, Messina V, Piro F, Tramontana L, Cherchi S, Falchi M, Bertuccini L, Spano F. 2022. Functional Characterization of the Thrombospondin-Related Paralogous Proteins Rhoptry Discharge Factors 1 and 2 Unveils Phenotypic Plasticity in Toxoplasma gondii Rhoptry Exocytosis. Front Microbiol 13:899243.

